# Uncoordinated centrosome duplication cycle underlies the instability of non-diploid states in mammalian somatic cells

**DOI:** 10.1101/194746

**Authors:** Kan Yaguchi, Ryo Matsui, Takahiro Yamamoto, Yuki Tsukada, Atsuko Shibanuma, Keiko Kamimura, Toshiaki Koda, Ryota Uehara

## Abstract

In animals, somatic cells are usually diploid and are unstable when haploid for unknown reasons. In this study, by comparing isogenic human cell lines with different ploidies, we found frequent centrosome loss specifically in the haploid state, which profoundly contributed to haploid instability through monopolar spindle formation and subsequent mitotic defects. We also found that efficiency of centriole licensing and duplication, but not that of DNA replication, changes proportionally to ploidy level, causing gradual loss or frequent overduplication of centrioles in haploid and tetraploid cells, respectively. Centriole licensing efficiency seemed to be modulated by astral microtubules, whose development scaled with ploidy level, and artificial enhancement of aster formation in haploid cells restored centriole licensing efficiency to diploid levels. Haploid-specific centrosome loss was also observed in parthenogenetic mouse embryos. We propose that incompatibility between the centrosome duplication and DNA replication cycles arising from different scaling properties of these bioprocesses upon ploidy changes, underlies the instability of non-diploid somatic cells in mammals.

**Summary:** Yaguchi et al. show that a delay or acceleration of centriole licensing compromises the control of centrosome number in haploid or tetraploid human cells, respectively, suggesting a cellular basis of the instability of non-diploid somatic cells in mammals.

## Introduction

Animal species generally have diplontic life cycles, where somatic cell division occurs only during the diploid phase. Exceptionally, haploid or near-haploid animal somatic cells arise through activation of oocytes in the absence of fertilization (e.g. parthenogenesis) or because of aberrant chromosome loss during tumorigenesis (Wutz, 2014). However, haploidy in animal somatic cells is generally unstable, and haploid cells in a wide variety of species, including insects, amphibians, and mammals, convert to diploid through doubling of the whole genome during successive culture for several weeks both in vitro and in vivo (Debec, 1984; Elling et al., 2011; Essletzbichler et al., 2014; Freed, 1962; Kaufman, 1978; Kotecki et al., 1999; Leeb and Wutz, 2011; Li et al., 2014; Sagi et al., 2016; Yang et al., 2013). This is in sharp contrast to plants and lower eukaryotic organisms, in which haploid somatic cells can proliferate stably (Forster et al., 2007; Mable and Otto, 1998). This raises the possibility that, specifically in animals, the cell replication mechanism is stringently adapted to the diploid state, and becomes compromised in haploid cells; however, the physiological impacts of ploidy differences on animal cell replication processes remain largely unknown.

In animal cells, control of centrosome number is essential for precise cell replication. During mitosis, pairs of centrosomes serve as major microtubule organizing centers (MTOCs) for bipolar spindle formation, and irregular numbers of centrosomes form spindles with abnormal polarities, endangering proper chromosome segregation (Gonczy, 2015). Centrosome number control is achieved through elaborate regulation of the centrosome duplication cycle. Upon exit from mitosis, an engaged pair of centrioles composing a centrosome separate from one another, producing two centrosomes (Kuriyama and Borisy, 1981). This centriole disengagement process is a prerequisite for “licensing” each pre-existing centriole to serve as a template for the formation of a daughter centriole in the subsequent cell cycle (Tsou and Stearns, 2006; Tsou et al., 2009). A scaffold protein, Cep152, accumulates on the licensed pre-existing centrioles, subsequently recruiting a key centriole duplication regulator, Polo-like kinase 4 (Plk4) (Cizmecioglu et al., 2010; Dzhindzhev et al., 2010; Fu et al., 2016; Hatch et al., 2010; Kim et al., 2013; Sonnen et al., 2013). Plk4, in turn, mediates the recruitment of SAS-6 on the outside wall of the pre-existing centrioles to form the procentriolar cartwheel, which founds a basis for the subsequent elongation of daughter centrioles (Bettencourt-Dias et al., 2005; Dzhindzhev et al., 2014; Fong et al., 2014; Habedanck et al., 2005; Kleylein-Sohn et al., 2007; Leidel et al., 2005; Moyer et al., 2015; Nakazawa et al., 2007; Ohta et al., 2014). Importantly, there are striking similarities between the molecular mechanisms governing temporal regulation of the centriole duplication cycle and DNA replication cycle. A mitotic kinase, Polo-like kinase 1 (Plk1), and a cysteine endoprotease, separase, cooperatively regulate resolution of the connections of the engaged centrioles or paired sister chromatids during or at the end of mitosis, and cyclin E-cdk2 controls the initiation of both centriole duplication and DNA replication during the G1/S phase (Coverley et al., 2002; Matsumoto et al., 1999; Meraldi et al., 1999; Nasmyth, 2002; Sumara et al., 2002; Tsou and Stearns, 2006; Tsou et al., 2009). These regulatory mechanisms ensure precise temporal coordination between these two cellular processes, allowing cells to possess a constant number of centrosomes throughout numerous rounds of cell cycles during proliferation.

To determine the cellular processes affected by ploidy difference and understand the origin of intolerance of somatic haploidy in animal cells, we carried out side-by-side comparisons of cell replication in isogenic mammalian somatic cells with different ploidy levels. We found that the efficiency of the centrosome cycle progression scales proportionally with ploidy level, which uncouples the progression of centrosome cycle from that of DNA cycle and compromises centrosome number control in non-diploid states.

## Results

### Haploid state-specific defects in mitotic progression in human somatic cells

To investigate the effect of differences in ploidy on the cell replication process, we used the near-haploid human cell line, HAP1 (Carette et al., 2011). As previously reported, the haploid state of this cell line was unstable and almost all cells in haploid-enriched culture diploidized over several weeks of passage (Fig. 1A) (Essletzbichler et al., 2014). Diploidized cells were significantly larger than haploid cells (Fig. 1B and S1A–C); therefore, we could purify the isogenic haploid and diploid cell populations separately for side-by-side comparisons, by size-based sorting.

**Fig. 1:**
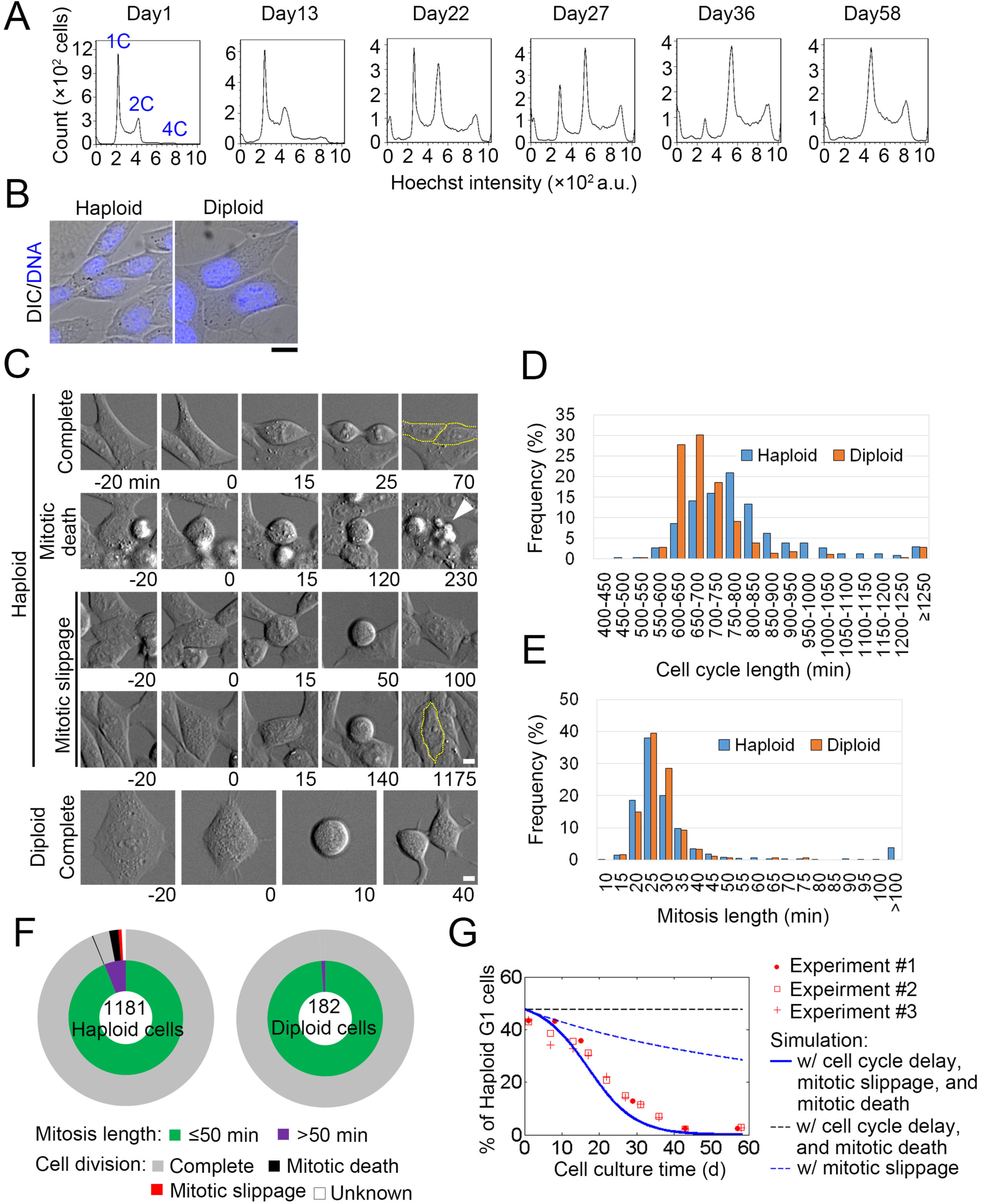
Haploid-specific mitotic defects and subsequent diploidization in HAP1 cells. (**A**) Flow cytometric analysis of DNA content in Hoechst-stained cells during long-term culture. (**B**) Hoechst-stained haploid and diploid cells. (**C**) Live images of haploid and diploid cells in mitosis taken at 5-min intervals. Nuclear envelope breakdown (NEBD) was set as 0 min. Arrowhead: debris from a cell that has undergone mitotic death. Broken lines indicate cell boundaries. Scale bars, 10 μm in B, and 5 μm in C. (**D**, **E**) Distribution of cell cycle length (from NEBD to the following NEBD), or mitotic duration (from NEBD to anaphase onset) quantified from 339 haploid and 285 diploid (D), and 1180 haploid and 182 diploid cells (E), respectively. Data are from 2 independent experiments. (**F**) Classification of mitotic defects (outer circle) sorted by mitotic duration (inner circle) determined based on analysis of 1181 haploid and 182 diploid cells in 2 independent experiments. (**G**) Time course of the haploid G1 fraction during long-term culture of haploid-enriched cells. Data from 3 independent experiments are compared with theoretical model simulations.

We first compared the progression of the cell cycle and cell division in haploid and diploid cells by live cell imaging (Fig. 1C–F, Video 1–3). Average cell cycle length was significantly greater in haploid than in diploid cells (806 ± 212 and 714 ± 186 min, respectively; n > 282; *p* < 10^-8^, *t*-test) (Fig. 1D). We also found that 72 of 1181 haploid cells (6.0%) exhibited severe mitotic delay, spending > 50 min in the mitotic phase (Fig. 1C, E, and F). Of mitotically arrested haploid cells, 36 entered anaphase and completed cytokinesis, while 20 died during mitosis, and the remaining 7 exited the mitotic phase without chromosome segregation (mitotic slippage; Fig. 1C and F, Video 2). Importantly, these mitotic defects were scarcely observed in diploid cells, suggesting that they were due to haploid-specific issues (Fig. 1F, Video 3).

Since mitotic slippage doubles DNA content, it can affect the stability of the haploid state, even when it occurs at low frequency. We therefore estimated the potential contribution of haploid-specific mitotic defects to haploid instability, using a mathematical cell population transition model (see *Materials and methods*). In this model, haploid or diploid cells proliferate exponentially, with respective doubling times corresponding to measured cell cycle lengths. Haploid cells die or diploidize through mitotic death or mitotic slippage, respectively, with empirically observed frequencies (Fig. 1F). A simulated time course of a haploid population using the model was essentially consistent with the experimentally observed diploidization profiles (Fig. 1G). In the simulation, mitotic slippage at the observed order of frequency could account solely for approximately 40% of haploid population loss during 60 days of culture, and haploid stability was sensitive to small changes in the frequency of mitotic slippage (Fig. 1G, and S1D). Based on the above experimental and theoretical analyses, we propose the following mechanism for haploid instability: haploid-specific cell division failure drives chronic conversion of haploid cells into diploid, which is accompanied by preferential expansion of the diploid population due to its cell-autonomous growth advantage over haploid cells. As haploid-specific mitotic defects are likely to contribute profoundly to haploid instability, we decided to investigate the cellular mechanisms underlying them.

### Frequent monopolar spindle formation in haploid human somatic cells

To determine the cause of haploid-specific mitotic defects, we investigated the organization of the mitotic spindle in haploid and diploid cells by α-tubulin immunostaining. Whereas the majority of diploid cells possessed bipolar spindles, ∼25% (n = 3) of haploid cells had monopolar spindles (Fig. 2A and B). Consistently, whereas the majority of diploid mitotic cells had a pair of centrosomes, each of which consisted of two centrioles, we observed loss of centrosomes and centrioles in > 20% of haploid mitotic cells (visualized by γ-tubulin and centrin immunostaining, respectively; Fig. 2C, D, and S1E). Since these haploid and diploid cells were isogenic, the frequent centrosome loss in haploid cells must be a consequence of haploidy rather than genetic background.

**Fig. 2:**
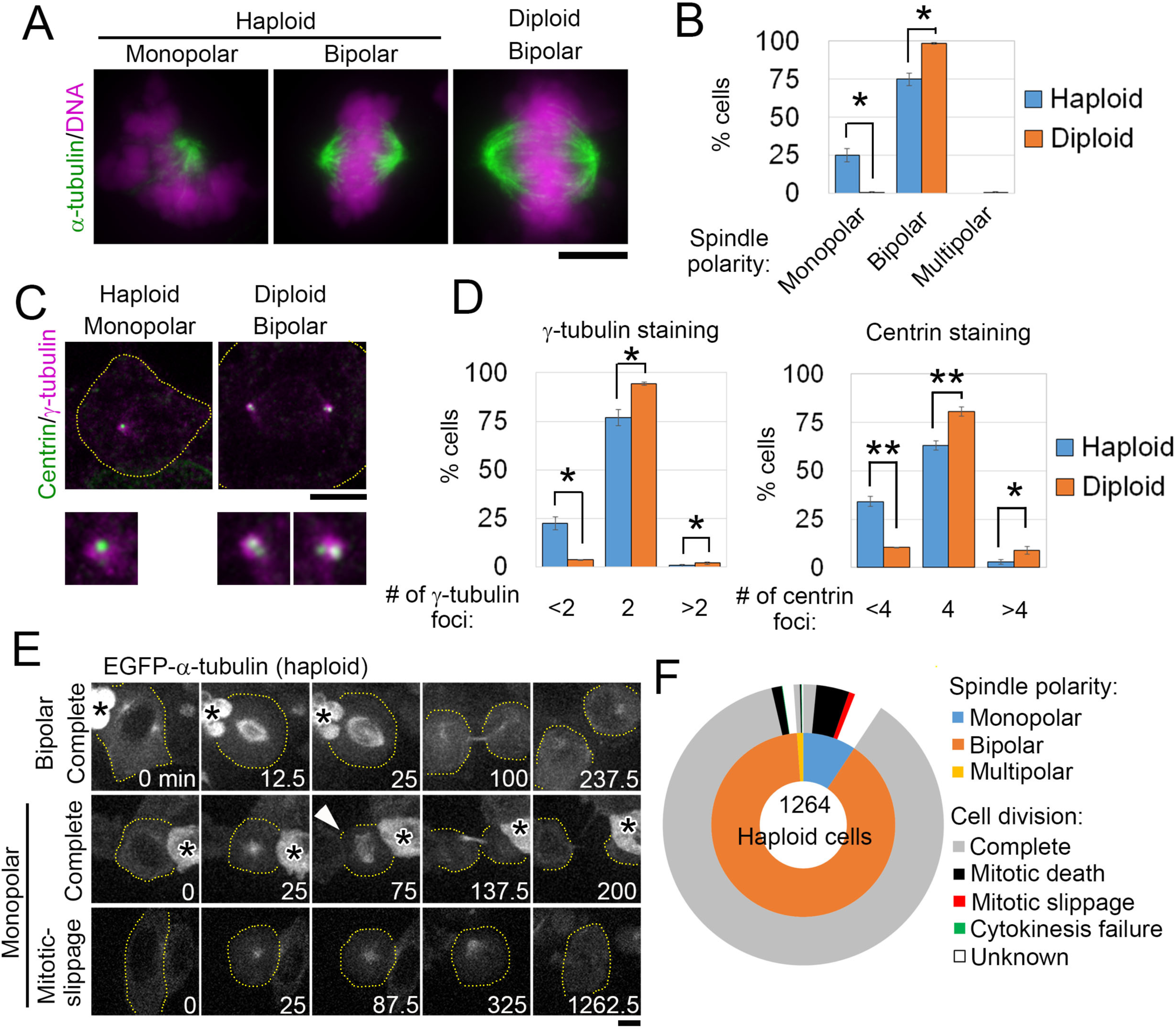
Centrosome loss and monopolar spindle formation in haploid HAP1 cells. (**A**, **C**) Immunostaining of α-tubulin and chromosomes (stained using DAPI) (A), or γ-tubulin and centrin (C) in haploid and diploid mitotic cells. Enlarged images (×3) of centrosomes are shown at bottom in C. (**B**, **D**) Frequency of spindle polarities (B) and centrosome or centriole numbers (D) in A and C, respectively. Means ± standard error (SE) of 3 independent experiments (* p < 0.05, ** p < 0.01, *t*-test). At least 159 (B), or 302 cells (D) were analyzed per condition. (**E**) Live images of haploid EGFP-α-tubulin cells taken at 12.5-min intervals. NEBD was set as 0 min. Arrowhead: an acentrosomal pole newly formed from a monopolar spindle. Broken lines indicate cell boundaries. Asterisks indicate neighboring cells. Scale bars, 5 μm. (**F**) Classification of mitotic defects (outer circle) sorted by spindle polarities (inner circle) determined by analysis of 1264 haploid EGFP-α-tubulin cells from 5 independent experiments. Cells that moved out of the field of view during the mitotic phase were categorized as unknown.

To examine the relationship between spindle disorganization and mitotic defects in haploid cells, we carried out live imaging using haploid HAP1 cells stably expressing EGFP-α-tubulin (Fig. 2E and F, Fig. S1F, and Video 4–6). Of 1264 cells that entered the mitotic phase, 118 (9.3%) formed monopolar spindles and of these monopolar cells, 19 later became bipolar through the formation of presumptive acentrosomal poles, and completed mitosis (arrowhead in Fig. 2E, and Video 5), 48 died in mitosis, and 9 underwent mitotic slippage (Fig. 2E, F, and Video 6). Importantly, all mitotic slippage events and the majority of mitotic deaths accompanied spindle monopolarization. These results suggest that spindle disorganization due to haploid-specific centrosome loss makes a major contribution to the mitotic defects observed in haploid cells.

### Centriole duplication efficiency scales proportionally with ploidy level

To understand how haploid cells lose their centrosomes, we next tested the progression of centriole duplication in haploid and diploid cells. Cells were synchronized by mitotic shake-off after release from nocodazole arrest, and DNA replication and centriole duplication in the subsequent cell cycle was monitored by BrdU incorporation and counting of immunostained centrin-positive foci, respectively (Fig. 3A–C, and S2A). BrdU incorporation in haploid cells was slightly slower than that in diploids; however, it reached the same maximum level within 8 h of nocodazole release (Fig. 3A). In contrast, the progression of centriole duplication was drastically delayed in haploid cells relative to that in diploids. Centriole duplication started around 5 h after nocodazole release and cells with four centrioles predominated at 8 h after nocodazole release in diploid cells, whereas ∼65% (n = 3) of haploid cells had unduplicated centrioles at that time (Fig. 3B and C). When the duration of the S phase was extended by treating cells with thymidine after nocodazole release, the percentage of duplicated pairs of centrioles in haploid cells increased monotonically, reaching a maximum level equivalent to that in diploids within 16 h after addition of thymidine (Fig. S2B). Hence, the centriole duplication process was not completely compromised in haploids, rather it became significantly less efficient and unable to keep pace with the DNA replication cycle, resulting in gradual loss of centrioles in successive cell cycles.

**Fig. 3:**
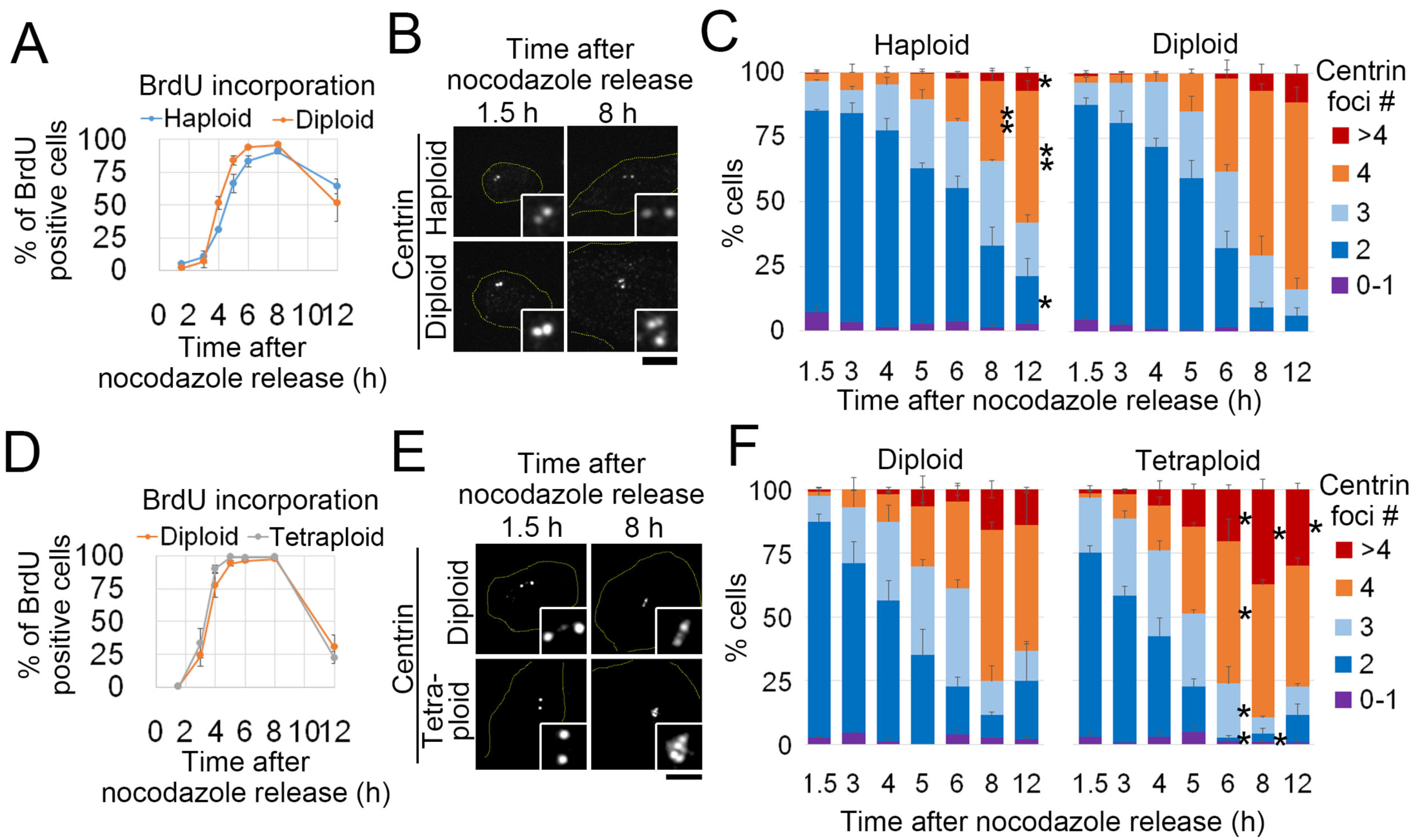
Centriole duplication efficiency scales to ploidy level in HAP1 cells. (**A**, **D**) BrdU incorporation after nocodazole release in cells with different ploidies. Mean ± SE of 3 independent experiments. At least 1258 (A) or 818 cells (D) were analyzed for each data point. (**B**, **E**) Immunostaining of centrin in synchronized cells with different ploidies. Scale bars, 5 μm. (**C**, **F**) Percentages of cells with indicated numbers of centrin foci at each time point after nocodazole release. Means ± SE of 3 independent experiments (asterisks indicate significant differences from diploid cells; * p < 0.05, ** p < 0.01, *t*-test). At least 170 (C), or 87 cells (F) were analyzed for each data point. Note that centrin foci number decreased from 8 h to 12 h after nocodazole release in some cases, because some population of cells divided during that time.

Next, we determined whether an increase in ploidy from diploid could also affect centriole duplication efficiency. We established stable tetraploid HAP1 cell lines by doubling the whole genomes of diploid cells (Fig. S2C; see *Materials and methods*). Tetraploid cells had an average of two mother centrioles when synchronized at the G1 phase, suggesting that excess centrosomes obtained upon tetraploidization had been lost during subsequent cloning, as reported for other cell lines (Fig. 3D–F) (Ganem et al., 2009; Potapova et al., 2016). The progression of BrdU incorporation after nocodazole release was similar between diploid and tetraploid cells (Fig. 3D and S2D); however, centriole duplication progressed significantly faster in tetraploids than that in diploids (Fig. 3E and F). We observed centriole overduplication significantly more frequently in tetraploid than in diploid cells (Fig. 3F). Therefore, the efficiency of centriole duplication scaled proportionally with ploidy level, whereas that of DNA replication was relatively insensitive to ploidy. This difference in ploidy dependency potentially threatens centrosome homeostasis upon ploidy conversion (either decreasing or increasing) from diploidy, owing to lack of coordination of these biological processes.

Next, we addressed the molecular mechanism by which ploidy difference affects centriole duplication efficiency. For this, we analyzed recruitment of the key duplication factors Cep152, Plk4, and SAS-6 on the pre-existing centrioles in haploid, diploid, or tetraploid cells synchronized at the G1/S phase by nocodazole release and mitotic shake-off (Fig. 4). At early G1 phase (2 h after nocodazole release), Cep152 had already accumulated to more than 50% (n = 3) of pre-existing centrioles in all ploidy states, which is consistent with previous reports that mother centrioles recruit Cep152 from the previous cell cycle (Fig. 4A and B) (Fu et al., 2016; Park et al., 2014; Sonnen et al., 2013). In diploid cells, the majority of pre-existing centrioles recruited Cep152 before entry into S phase, but recruitment of Cep152 was drastically delayed in haploid cells, and ∼35% (n = 3) of centrioles remained devoid of Cep152 signals even in early S phase (4 h after nocodazole release) (Fig. 4B). In contrast, Cep152 recruitment was accelerated in tetraploid cells compared to that in diploid cells. In haploid or tetraploid cells, subsequent recruitments of Plk4 and SAS6 also fell behind or were ahead of, respectively, those in diploid cells, which is consistent with their hierarchical localization dependencies on Cep152 (Kim et al., 2013; Sonnen et al., 2013) (Fig. 4C–F). The time gaps of the recruitment of these proteins among different ploidy states roughly corresponded with those of daughter centriole formation (Fig. 3C and F), suggesting that the ploidy-linked change in centriole duplication efficiency arises from the change in the timing of the recruitment of these key duplication factors. Besides the changes in the timing of their recruitment, we also found that amounts of Cep152 and Plk4 recruited to the centrioles during the S phase (6 h after nocodazole release) changed proportionally with ploidy levels (Fig. S2E and F).

**Fig. 4:**
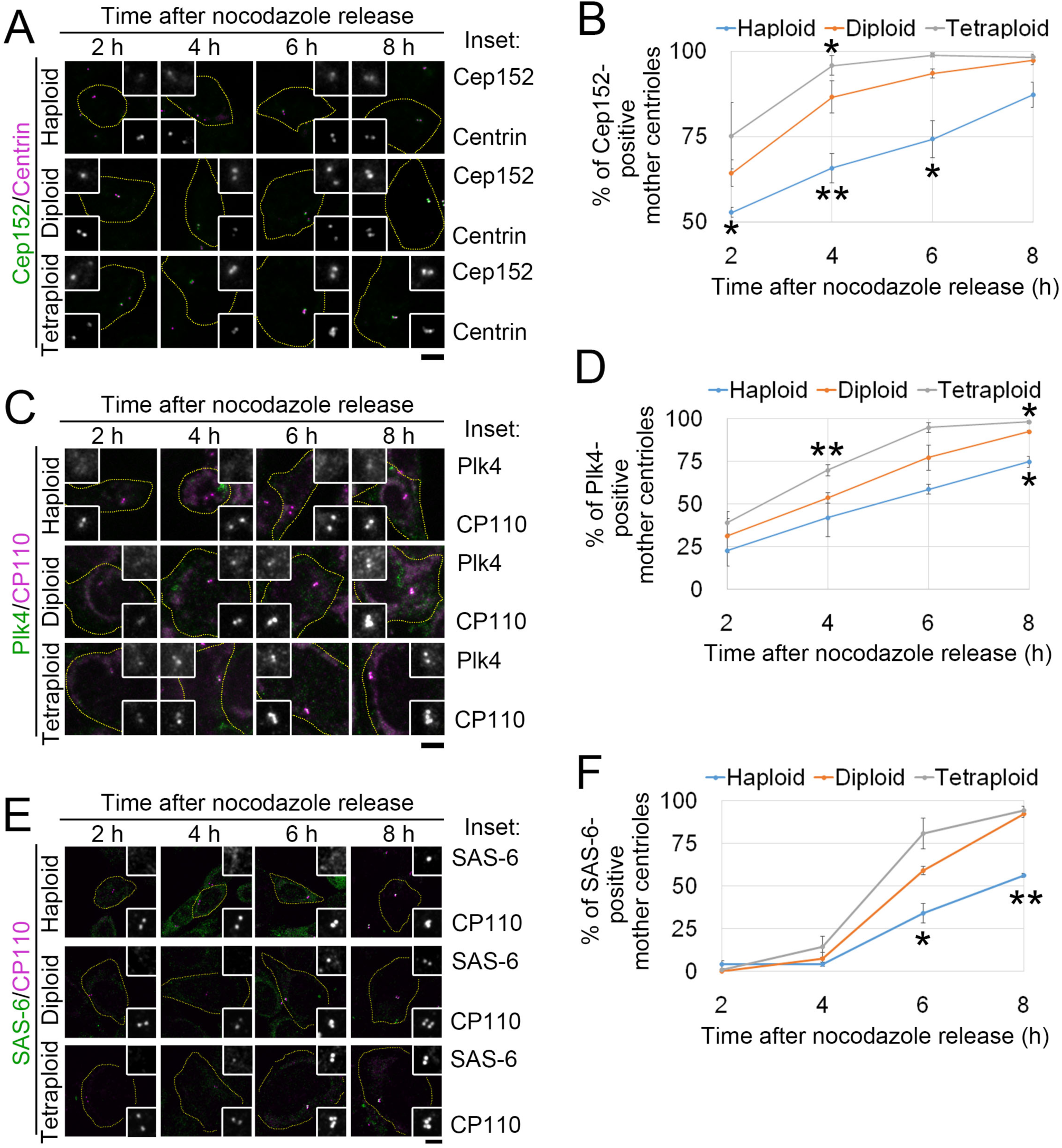
Recruitment efficiency of centriole duplication factors scales to ploidy level. (**A**, **C**, **E**) Immunostaining of Cep152 (A), Plk4 (C), and SAS-6 (E) in synchronized cells with different ploidies. The centrioles were marked by immunostaining of centrin (A) or CP110 (C and E). Broken lines indicate cell boundaries. Insets: ×3 enlarged images of the centrioles. Scale bars, 5 μm. (**B**, **D**, **F**) Percentages of Cep152-, Plk4-, and SAS-6-positive mother centrioles in A, C, or E. Means ± SE of 3 independent experiments (asterisks indicate significant differences from diploid cells; * p < 0.05, ** p < 0.01, *t*-test). At least 114 (B), 147 (D), or 250 mother centrioles (F) were analyzed for each data point.

### Centriole licensing is rate limiting for the centriole duplication process in different ploidies

During mitotic exit, the tightly connected mother and daughter centrioles are disengaged, which releases the intrinsic blocks for duplication and “licenses” these centrioles to recruit key centriole duplication factors and serve as templates for centriole biogenesis (Tsou and Stearns, 2006; Tsou et al., 2009). Because the centriole licensing through disengagement may potentially be rate limiting for the entire centriole duplication process, we next investigated the effect of ploidy difference on the status of centriole engagement at the mitotic exit. For that purpose, a centriole-linking protein, C-Nap1, and centrin were co-immunostained in asynchronous haploid, diploid, and tetraploid cells (Fig. 5A and B). Using these markers, engaged centriole pairs are visualized as two adjacent centrin dots flanking a C-Nap1 dot, while disengaged pairs as two centrin dots with two discrete C-Nap1 dots (Tsou and Stearns, 2006). At mitotic exit, during which two daughter cells were connected by an intercellular bridge after the constriction of the contractile ring, ∼25% (n = 3) of diploid cells possessed a disengaged centriole pair (Fig. 5B). In haploids or tetraploid cells, frequency of the cells with disengaged centrioles was significantly lower or higher than that in diploids, respectively (Fig. 5B), suggesting that the efficiency of centriole disengagement scales to ploidy level.

Next, to understand the time course of the resolution of inter-centriolar connection during mitotic exit, we performed live imaging of haploid, diploid, and tetraploid cells stably expressing GFP-tagged centrin as a centriole marker (Fig. 5C–E, and S1F). With the spatiotemporal resolution of our live imaging setting, it was difficult to unambiguously specify the exact timing of centriole disengagement. However, by quantifying the time course of inter-centriolar distance during mitotic exit, we were able to follow the process of centriole separation (Fig. 5D and E). In all ploidy states, pairs of centrioles were located within 0.6 μm of one another at cytokinesis onset (Fig. 5D), which approximately corresponds with the inter-centriolar distance of engaged centriole pairs (Piel et al., 2000). By 30 min after cytokinesis onset, in 50% of diploid cells (6 out of 12 cells), inter-centriolar distance became more than 0.8 μm (Fig. 5E), which approximately corresponds to that of disengaged centriole pairs (Piel et al., 2000). In haploid cells, the progression of centriole separation was severely delayed (Fig. 5D), and inter-centriolar distance stayed below 0.8 μm in more than 30% of them (4 out of 13 cells), even at 140 min after cytokinesis onset (Fig. 5E). In contrast, in tetraploid cells, the timing of the centriole separation was substantially brought forward compared to that in diploid cells (Fig. 5D and E). Therefore, consistent with the result from the fixed cells analysis, the efficiency of centriole separation during mitotic exit scaled with ploidy level.

**Fig. 5:**
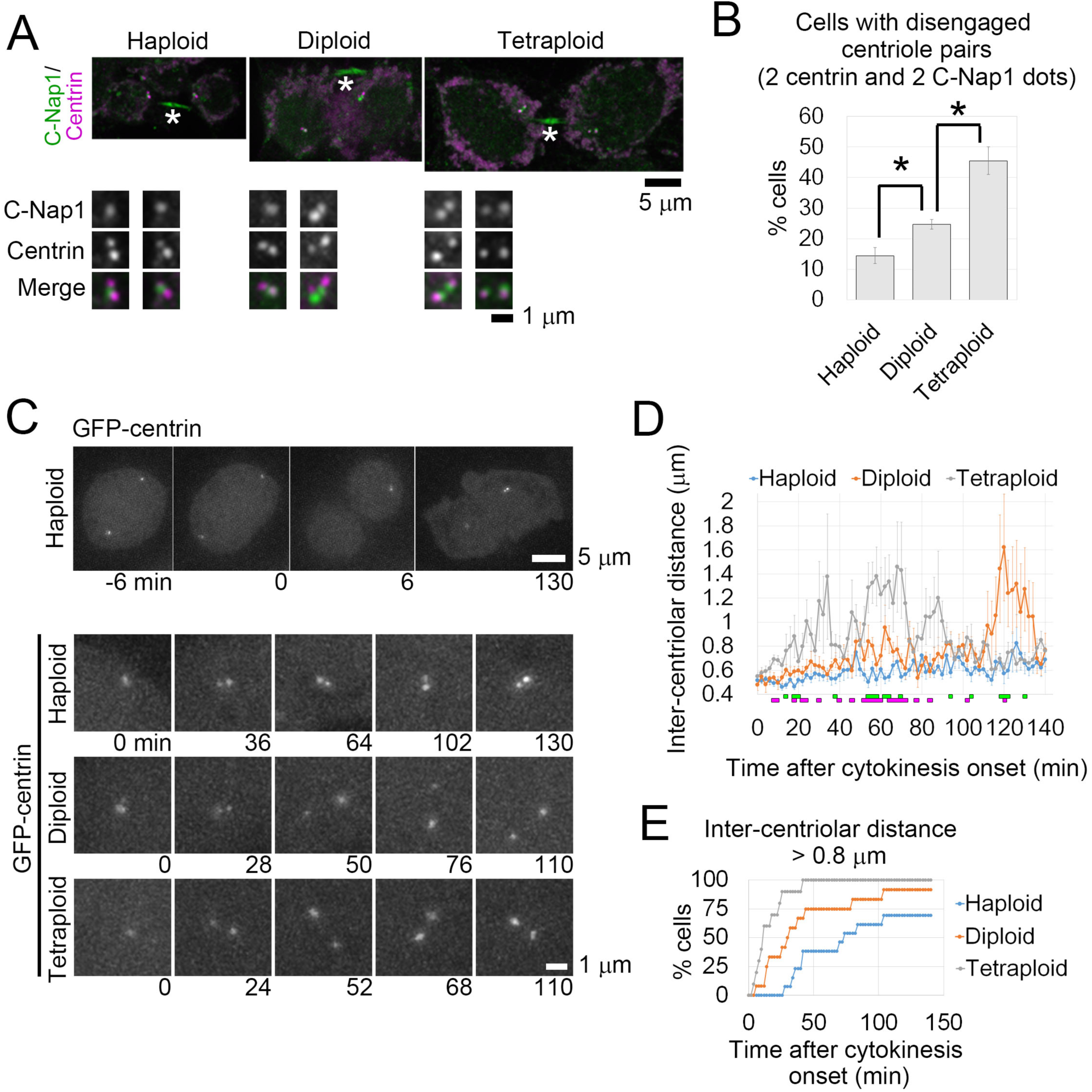
The efficiency of centriole disengagement scales to ploidy level. (**A**) Immunostaining of centrin and C-Nap1 in haploid, diploid, and tetraploid cells at mitotic exit. Whole cell images (top) and ×3 enlarged images of the centrioles (bottom). Asterisks indicate non-specific staining of the intercellular bridge. (**B**) Frequencies of cells with disengaged centriole pairs (with 2 centrin and 2 C-Nap1 dots) in A. Means ± SE of 3 independent experiments (* p < 0.05, *t*-test). At least 144 cells were analyzed for each condition. (**C**) Live imaging of GFP-centrin cells taken at 2-min intervals. Cytokinesis onset was set as 0 min. Whole cell images of a haploid cell (top) and enlarged images of the centrioles in cells with different ploidies (bottom). (**D**) Time course of inter-centriolar distance in C. Means ± SE of at least 9 cells from at least 2 independent experiments for each data point (at least 10 cells were analyzed for each condition). Green or magenta markers at the bottom of the graph indicate statistically significant differences between haploid and diploid or diploid and tetraploid cells, respectively (p < 0.05, *t*-test). (**E**) Cumulative frequency of cells in which inter-centriolar distance had reached 0.8 μm in C. At least 10 cells from at least 2 independent experiments were analyzed for each condition.

To assess whether the ploidy-dependent change in the efficiency of centriole disengagement is the primary cause for the delay in the progression of centriole duplication in haploid cells, we set out to manipulate the efficiency of centriole disengagement. Previous studies have revealed that a particular fraction of pericentriolar material (PCM) is involved in centriole engagement, and that the removal of a PCM component, PCNT/pericentrin/kendrin, from the centrosomes is the prerequisite for timely disengagement of centriole pairs and subsequent centriole duplication (Lee and Rhee, 2012; Matsuo et al., 2012; Pagan et al., 2015). Therefore, we investigated the effect of PCNT depletion on the progression of centriole separation during mitotic exit in GFP-centrin-expressing haploid cells by live imaging (Fig. 6). To avoid the potential side effects of the prolonged mitotic phase caused by PCNT depletion (Zimmerman et al., 2004), we co-depleted mad2, which is essential for activation of the spindle assembly checkpoint (Chen et al., 1996; Waters et al., 1998). Live imaging was conducted 48 h after RNAi treatment, when distribution of DNA content either in mad2-depleted or PCNT- and mad2-co-depleted haploid cells remained unchanged from that in mock-depleted haploid control cells (Fig. 6A and B). To avoid complexity of interpretation caused by cell division defects or failure, we analyzed only the cells that succeeded in symmetric cell division without mitotic arrest during live imaging. The progression of centriole separation was drastically accelerated in PCNT- and mad2-co-depleted haploid cells compared to that in mock-depleted or mad2-depleted haploid control cells (Fig. 6C–E). Inter-centriolar distance became more than 0.8 μm by 30 min in more than 50% of PCNT- and mad2-co-depleted cells (6 out of 10 cells) (Fig. 6E). Therefore, forced removal of PCNT restored centriole separation efficiency in haploid cells to the diploid level.

**Fig. 6:**
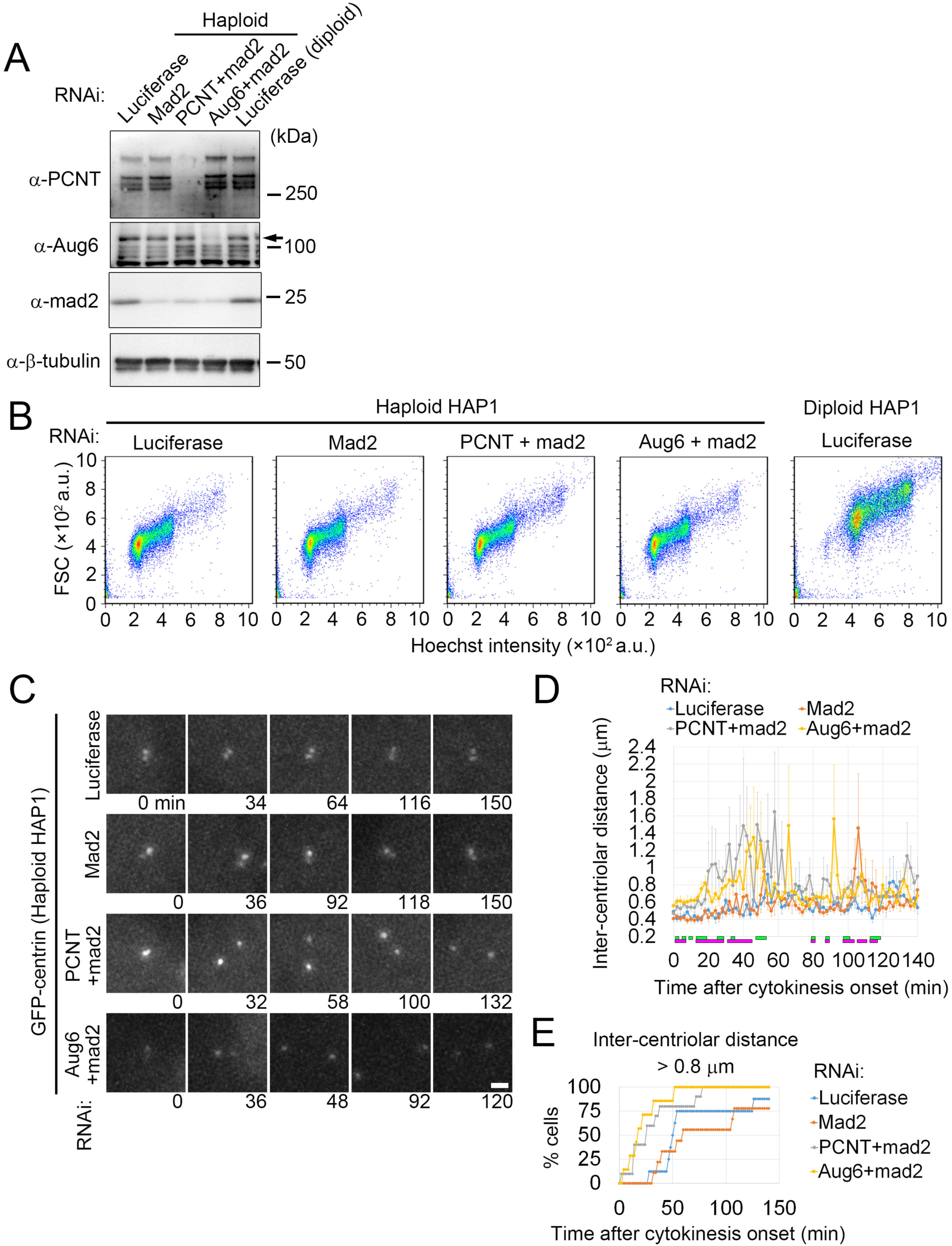
Depletion of PCNT or augmin accelerates centriole separation in haploid cells. (**A**) Immunoblotting of PCNT, Aug6, and mad2 in RNAi-treated haploid and diploid HAP1 cells. β-tubulin was detected as a loading control. The arrow indicates specific bands of Aug6. (**B**) Flow cytometric analysis of FSC and DNA content in RNAi-treated haploid and diploid HAP1 cells stained by Hoechst. (**C**) Live imaging of RNAi-treated haploid GFP-centrin cells taken at 2-min intervals. Cytokinesis onset was set as 0 min. Enlarged images of the centrioles are shown. Scale bar, 1 μm. (**D**) Time course of inter-centriolar distance in C. Means ± SE of at least 5 cells from 2 independent experiments for each data point (at least 7 cells were analyzed for each condition). Green or magenta markers at the bottom of the graph indicate statistically significant differences between mock-depleted and PCNT- and mad2-co-depleted, or Aug6- and mad2-co-depleted haploid cells, respectively (p < 0.05, *t*-test). (**E**) Cumulative frequency of cells in which inter-centriolar distance had reached 0.8 μm in C. At least 7 cells from 2 independent experiments were analyzed for each condition.

We next tested the effect of PCNT depletion on the subsequent progression of centriole duplication in haploid cells. For that purpose, we performed correlative live- and fixed-cell imaging. RNAi-treated cells that expressed GFP-centrin were first monitored by DIC time-lapse imaging for 12–14 h to identify the cells that passed the mitotic phase (Fig. 7A). Cells were then fixed and subjected to fluorescence microscopy to count GFP-centrin dots in the live-monitored cells that were identified by grid patterns on the culture dishes (Fig. 7A and B). By quantifying the distribution of centriole numbers in respective time windows after mitotic exit, we were able to monitor and compare the progression of centriole duplication among different RNAi conditions (Fig. 7C). Consistent with the result in the synchronization assay (Fig. 3C), the progression of centriole duplication in mock-depleted haploid cells was severely delayed compared to that in the mock-depleted diploid counterpart (Fig. 7C). However, PCNT- and mad2-co-depletion considerably accelerated the progression of centriole duplication in haploid cells to a level similar to that in mock-depleted diploid cells (Fig. 7C). These results show that the delay in centriole disengagement is the primary cause of the delay in the progression of centriole duplication in haploid cells.

**Fig. 7:**
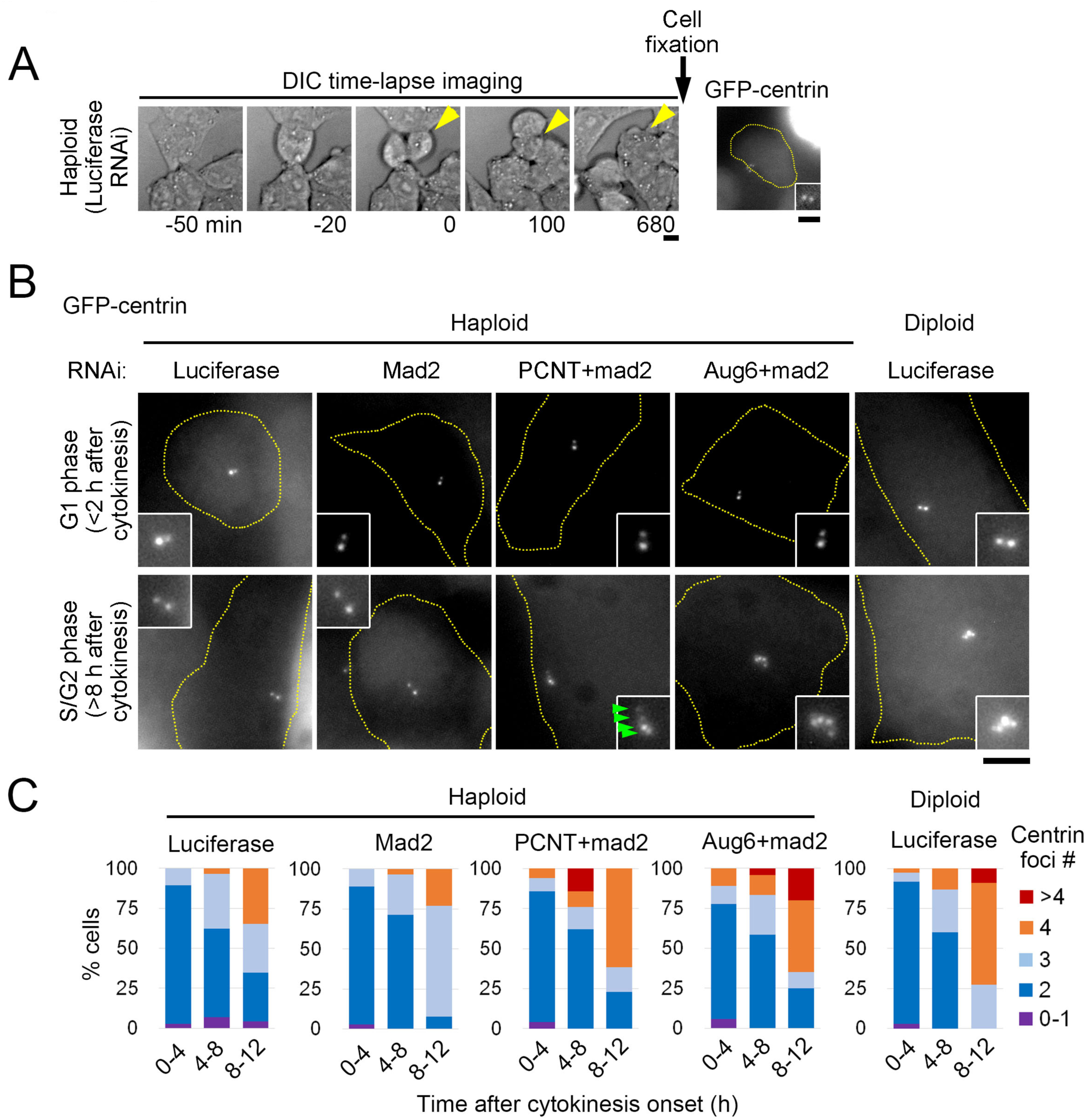
Depletion of PCNT or augmin accelerates centriole duplication in haploid cells. (**A**) Correlative live- and fixed-cell imaging for monitoring the progression of centriole duplication.RNAi-treated GFP-centrin cells were live-imaged by DIC microscopy every 10 min for 12–14 h. After fixation, the cells that passed the mitotic phase during time-lapse imaging were identified using grid patterns on the culture dish, and GFP-centrin in these cells was observed using fluorescence microscopy (yellow arrowheads and broken line). Insets: ×3 enlarged images of the centrioles. Cytokinesis onset was set as 0 min. (**B**) Fluorescence microscopy of GFP-centrin in RNAi-treated haploid and diploid HAP1 cells in the correlative live- and fixed-cell imaging. Centrin dots are indicated by arrowheads in a PCNT-depleted cell at the S/G2 phase. Insets: ×2 enlarged images of the centrioles. Scale bars, 5 μm. (**C**) Percentages of cells with indicated numbers of GFP-centrin foci at each time window after cytokinesis onset in B. At least 62 cells from at least 2 independent experiments were analyzed for each condition (>10 cells for each data point).

### Ploidy-dependent difference in astral MT development determines the efficiency of centriole licensing

We next wished to understand the mechanism that changes the efficiency of centriole disengagement in a ploidy-dependent manner. Previous studies have proposed two distinct mechanisms that drive disengagement of the centriole pair during mitotic exit, that is, the Plk1- and separase-mediated degradation of PCNT or mechanical separation of the engaged centriole pair by astral MT-driven pulling forces (Cabral et al., 2013; Kim et al., 2015; Lee and Rhee, 2012; Matsuo et al., 2012; Seo et al., 2015). Therefore, we tested whether either of these two mechanisms were affected by ploidy difference. We first tested the progression of PCNT degradation in haploid, diploid, and tetraploid cells that were arrested at and released from mitotic phase by nocodazole shake-off (Fig. 8). The degradation of PCNT analyzed by immunoblotting progressed in a similar manner in all ploidy states, suggesting that this process was unaffected by ploidy difference. We next assessed the possible effect of ploidy difference on astral MT-dependent centriole disengagement. Interestingly, immunostaining of α-tubulin revealed that during anaphase, in which centriole separation normally initiates, the cells with higher ploidies developed more prominent astral MTs than those with lower ploidies (Fig. 9A). Tracings of astral fibers revealed that the number of astral fibers associated with the cell cortex at the polar regions increased more than 2-fold upon the doubling of ploidies (Fig. 9B). Since astral MT generation is dependent on the MTOC at the spindle poles, we also compared accumulation of an essential MTOC factor, γ-tubulin, at the spindle poles among different ploidies (Fig. 9C and D). The accumulation of γ-tubulin scaled to ploidy levels, which is likely to promote the ploidy-dependent increase in astral MT generation.

**Fig. 8:**
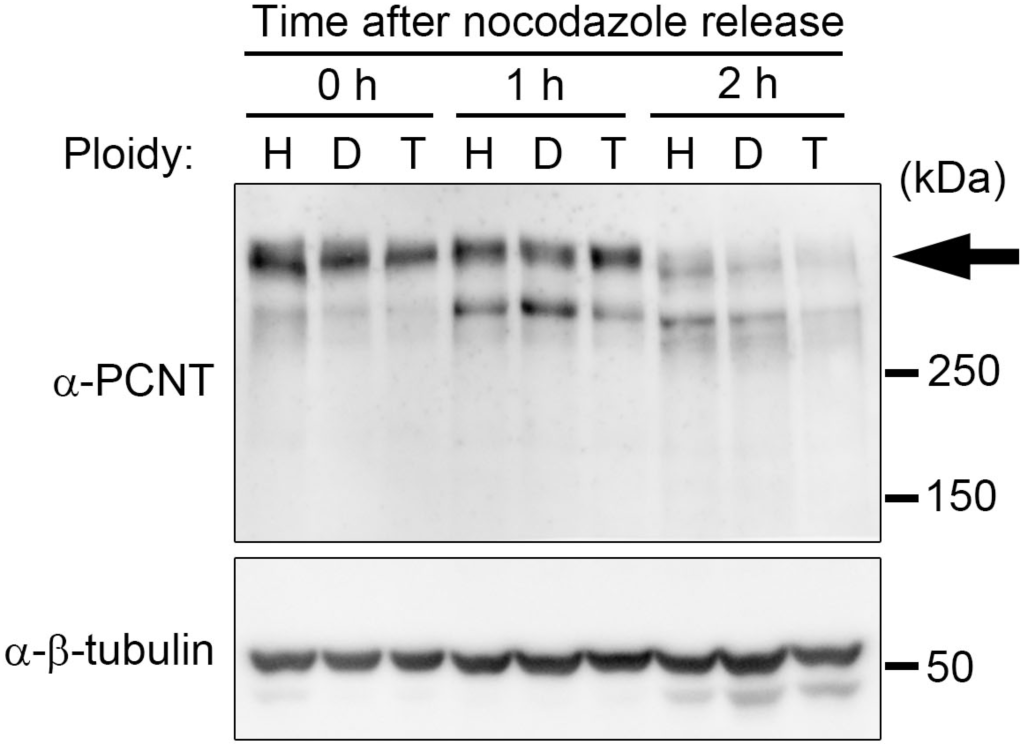
Progression of PCNT degradation is not affected by ploidy difference. Immunoblotting of PCNT before and after nocodazole release and mitotic shake-off in cells with different ploidies. β-tubulin was detected as a loading control. Arrow indicates intact PCNT.

**Fig. 9:**
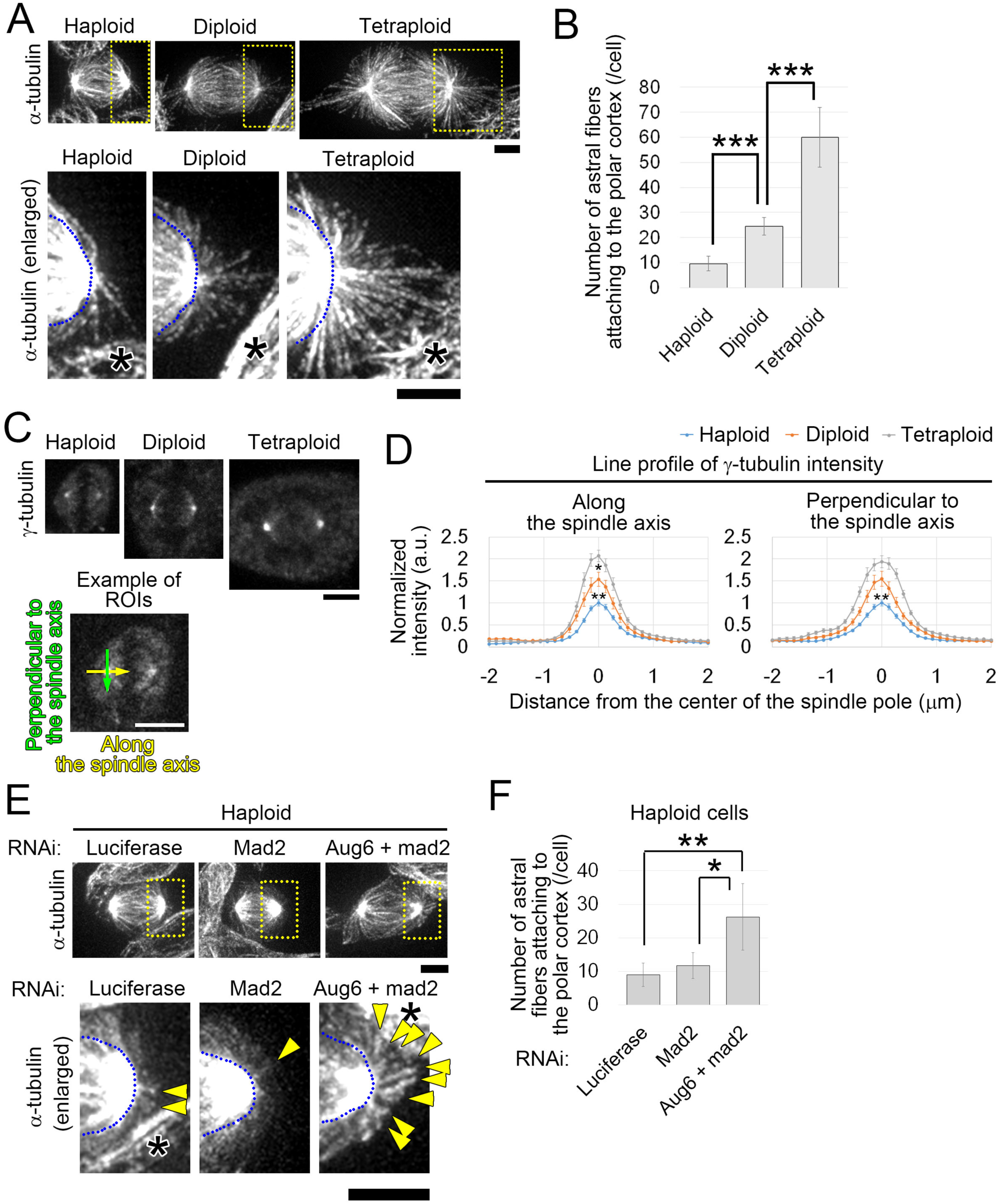
Development of astral MTs scales to ploidy level. (**A**, **E**) Immunostaining of α-tubulin during anaphase in cells with different ploidies (A) and RNAi-treated haploid cells (E). Enlarged images of the astral MTs at the polar region (yellow boxes) are shown in the bottom panels. Asterisks indicate neighboring interphase cells. Blue broken lines indicate spindle regions. Arrowheads in E indicate astral MTs. (**B**, **F**) Number of astral fibers attached to the polar cortex in A or E. Means ± SD of at least 5 cells (B) or at least 4 cells (F) from 2 independent experiments (* p < 0.05, ** p < 0.01, *** p < 10^-5^, *t*-test). Examples of MT tracking are shown in Video 7 or Video 8. (**C**) Immunostaining of γ-tubulin in pre-anaphase cells with different ploidies. Bottom: An example of regions of interest (ROIs) for the quantification in D. Scale bars, 5 μm. (**D**) Line profiles of γ-tubulin at the spindle poles in C. Means ± SE of at least 13 spindle poles in 8 cells from 2 independent experiments. Asterisks indicate statistically significant differences either between haploid and diploid or diploid and tetraploid cells at the center of the spindle pole (* p < 0.05, ** p < 0.01, *t*-test).

Next, to clarify the causal relationship between ploidy-dependent astral MT development and the efficiency of the centriole duplication cycle, we assessed whether artificial enhancement of astral MT formation can override inefficient centriole disengagement and duplication in haploid cells. Previously, we had found that depletion of a spindle-associated protein complex, augmin, suppressed MT generation within the mitotic spindle and instead enhanced development of prominent astral MTs during anaphase in HeLa cells (Uehara et al., 2016). Therefore, we tested the effect of augmin depletion on the progression of centriole disengagement and subsequent centriole duplication in haploid HAP1 cells. To avoid mitotic arrest caused by augmin depletion (Uehara et al., 2009), we co-depleted mad2 with an augmin subunit, Aug6 (Fig. 6A). Neither haploid DNA content nor cell size changed from mock-depleted control in Aug6- and mad2-co-depleted cells when experiments were conducted (Fig. 6B). In Aug6- and mad2-co-depleted haploid cells, the number of astral fibers associated with the polar cortex significantly increased compared to that in control haploid cells and became equivalent to that in diploid cells (Fig. 9E and F, see also Fig. 9B). Co-depletion of Aug6 and mad2 drastically accelerated the progression of centriole separation in haploid GFP-centrin2-expressing cells compared to that in control haploid cells (Fig. 6C–E), and in the correlative live- and fixed-cell imaging assay, efficiency of centriole duplication in Aug6- and mad2-depleted cells became similar to that in control diploid cells (Fig. 7B and C). The above results indicate that extent of astral MT development is a critical determinant of the efficiency of the centriole licensing process, and that the inefficient centriole duplication cycle in haploid cells is largely attributable to the poor organization of astral MTs during the mitotic phase.

### Haploid-specific centrosome loss in mouse parthenogenetic embryos

Finally, to assess whether the ploidy-linked change in centrosome homeostasis is also seen in other systems than human cultured cells, we investigated mitotic spindle organization in haploid and diploid mouse parthenogenetic embryos at stage E4.5, by which *de novo* centrosome generation has been completed (Courtois et al., 2012; Gueth-Hallonet et al., 1993; Latham et al., 2002; Liu et al., 2002) (see *Materials and methods*). As previously reported, morphological abnormalities were frequently observed in haploid embryos, whereas the majority of diploid embryos had normal morphology (Latham et al., 2002; Liu et al., 2002) (Fig. 10A, B, Fig. S3A, and B). A substantial proportion of mitotic cells in haploid embryos (46 out of 114 cells) had monopolar spindles with less than 2 centrosomes, whereas the majority of mitotic cells had bipolar spindles in diploid embryos (Fig. 10C–E). The haploid-specific centrosome loss was also observed in parthenogenetic embryos from another mouse strain (Fig. S3C and D). These results suggest conservation of the ploidy-centrosome link among mammalian organisms.

**Fig. 10:**
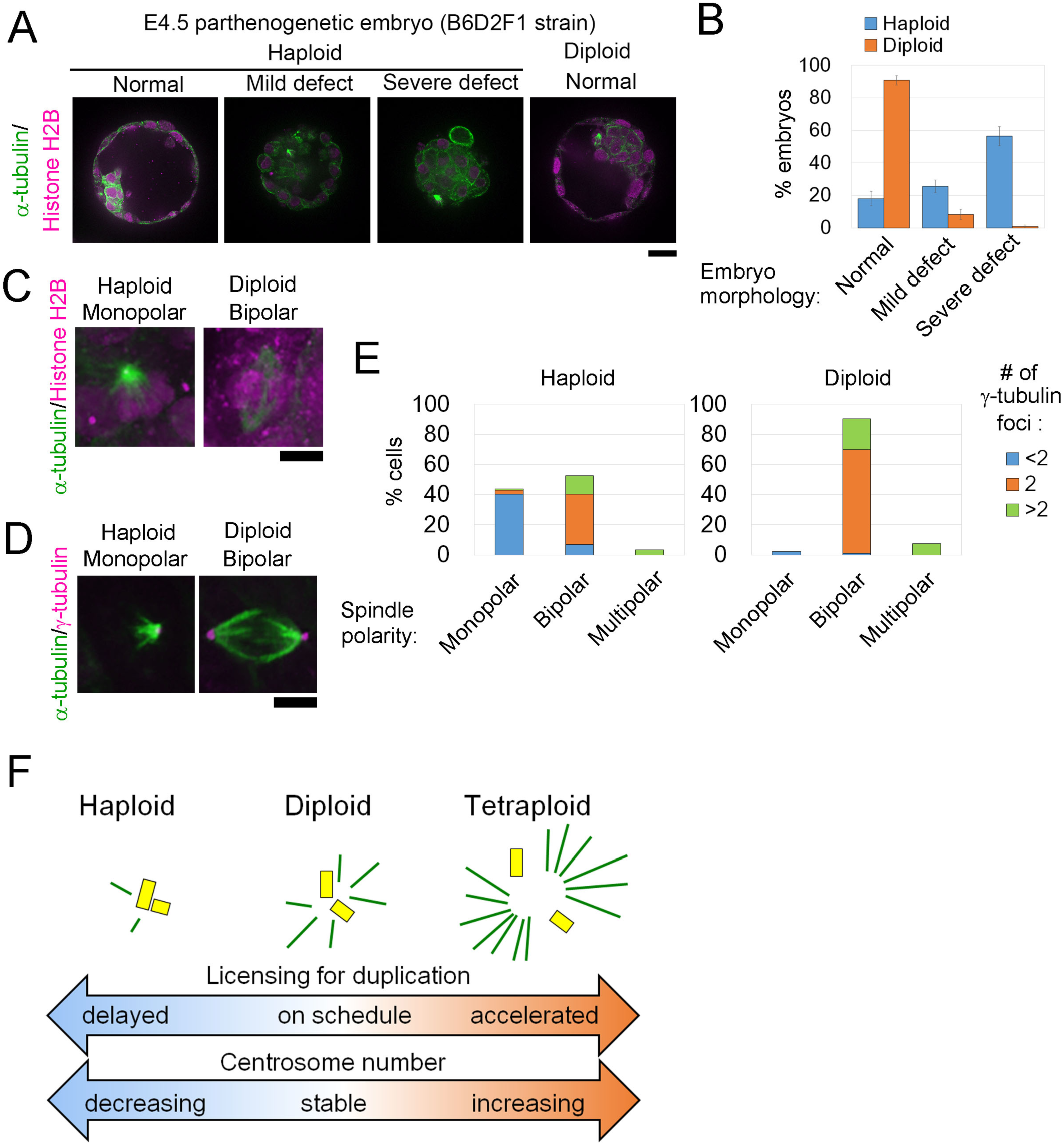
Centrosome reduction and monopolar spindle formation in haploid mouse embryos. (**A**, **C**, **D**) Immunostaining of α-tubulin and histone H2B (A and C), or γ-tubulin (D) in B6D2F1 mouse parthenogenetic embryos at stage E4.5. Mitotic cells are shown in C and D. Scale bars, 20 μm in A, and 5 μm in C and D. (**B**) Frequencies of morphological abnormalities in A. Means ± SE of 4 independent experiments. At least 109 embryos were analyzed per condition. (**E**) Spindle polarities and numbers of γ-tubulin foci in D. At least 93 cells from 71 embryos from 11 independent experiments were analyzed per condition. (**F**) A model for the ploidy-centrosome link. During mitosis, astral MTs (green lines) develop in a ploidy-linked manner. Inadequate or excess amounts of astral MTs lead to deceleration or acceleration of centriole disengagement (and subsequent duplication), respectively, in haploid or tetraploid cells, respectively. As a result, the efficiency of the entire centriole duplication cycle scales proportionally to ploidy level, which drives uncoupling of centriole duplication and DNA replication in non-diploid states.

## Discussion

The unstable nature of somatic haploidy has been recognized for decades in a wide variety of animal organisms, but the fundamental problems in maintaining the genomic integrity of somatic cells brought about by ploidy difference are poorly understood. Through side-by-side comparison of isogenic cell lines with different ploidy levels, we found a linear relationship between ploidy level and efficiency of the centrosome duplication cycle. This relationship seems to stem from the ploidy-linked scaling of astral MT development during mitosis, which promotes centriole disengagement in a ploidy-dependent manner (Fig. 10F). This ploidy-centrosome link impairs temporal coordination between the centrosome duplication cycle and the chromosome replication cycle in non-diploid states, which would account at least in part for the relatively low tolerance of animal cells to ploidy variance compared to acentrosomal organisms such as land plants.

### Ploidy-linked scaling of centrosomal protein accumulation and astral MT development

Our data indicate that astral MTs make a profound contribution to the centriole licensing process in mammalian cells as previously proposed in early *C. elegans* embryos (Cabral et al., 2013), and moreover, that development of astral MTs is one key determinant of the efficiency of the centriole duplication cycle in different ploidy states. Since γ-tubulin provides templates for astral MT nucleation from the centrosomes (Oakley et al., 2015), the ploidy-linked scaling of astral MT development would result from the ploidy-dependent accumulation of γ-tubulin at the spindle poles. The ploidy-dependent increase in centrosomal γ-tubulin seems to be a general consequence of the increased pool of centrosomal gene products available per centrosome scaffold upon ploidy increase. Similar ploidy-linked scaling of the centrosome equivalents has been observed in different fungal species (Storchova et al., 2006; Suzuki et al., 1982), suggesting generality of this phenomenon among a wide range of eukaryotic organisms.

Besides centrosomal protein accumulation at the spindle poles during mitosis, the accumulation of some key centriole duplication factors, such as Cep152 and Plk4, also changed in a ploidy-dependent manner during the G1/S phase. The general reduction in the amounts of centriole duplication factors at the centrosomes potentially endangers centrosome homeostasis in haploid states, especially considering that some important duplication factors such as Plk4 or NDC1 in yeast cells show haploinsufficiency (Chial et al., 1999; Ko et al., 2005). However, we found that the acceleration of centriole separation by PCNT or augmin depletion singly recovered the efficiency of centriole duplication in haploid cells to the diploid level. These results indicate that the delay in centriole licensing is the primary cause of the inefficient centriole duplication cycle in haploid cells, and that single gene copies of centriole duplication factors are sufficient to support the normal progression of centriole duplication in a haploid background. However, we cannot rule out the possibility that ploidy-dependent accumulation of centriole duplication factors influences the efficiency of centriole duplication in hyperploidy states. It is also important that future studies examine whether ploidy-dependent changes in centrosomal protein enrichment have any influence on the ultrastructure of the centriole throughout the centrosome duplication cycle.

### Generality of the ploidy-centrosome link among animal organisms

In addition to the loss in cultured haploid human cells, we observed centrosome loss in about 40% of mitotic cells in mouse parthenogenetic embryos, suggesting that the link between ploidy and centrosome cycle efficiency is preserved broadly in mammalian somatic cells. On the other hand, it remains to be elucidated whether a similar ploidy-centrosome link is conserved among non-mammalian animal species. Of interest, however, is that in different fish species, haploid embryos develop abnormally small brains and eyes (Araki et al., 2001; Luo and Li, 2003; Patton and Zon, 2001), which are very similar to morphological defects reported in zebrafish embryos depleted of several centrosomal genes (Kim et al., 2011; Novorol et al., 2013). Therefore, it is intriguing to speculate that these developmental defects observed in haploid fishes are due to haploidy-driven centrosome loss. Another hint on the generality of the ploidy-centrosome link is that haploid cell lines derived from haploid lethal embryos of a *Drosophila melanogaster* mutant are devoid of centrioles (Debec, 1978; Debec, 1984; Debec and Abbadie, 1989; Szollosi et al., 1986). Although in the original papers, the centrosome loss in these cell lines was attributed to an unidentified mutation in the original embryos, it is also possible that their haploid state *per se* drove the centrosome loss, as seen in the mammalian cells in this study. Further analyses on the mechanism of centrosome loss in these cell lines would improve our understanding of the general impact of ploidy difference on centrosome homeostasis in a wider variety of animals.

### Potential importance of the hyperploidy-driven centriole overduplication

Our finding of the hyperploidy-driven upregulation of centriole duplication suggests a new route to centrosome amplification in hyperploid tumor cells. Hyperploid tumor cells often possess extra centrosomes, which conceivably arise from cell division failure in ancestor cells (Godinho and Pellman, 2014). A prevailing view is that these extra centrosomes are subsequently lost in the majority of progeny cells, but are sustained in a small population, selected owing to their invasive qualities (Ganem et al., 2009; Godinho et al., 2014). The hyperploidy-driven centrosome overduplication identified in this study would promote chronic centrosome amplification without assuming any selective processes or genetic rearrangements of centriole duplication genes, and thus, may make profound contribution to the maintenance of extra centrosomes in hyperploid tumor cells. An intriguing future study would be to investigate the contribution of hyperploidy-driven centrosome overduplication to genome instability during cancer development using *in vivo* models.

## Materials and Methods

### Cell culture and flow cytometry

The HAP1 cell line was purchased from Haplogen GmbH (Vienna, Austria) and cultured in Iscove’s Modified Dulbecco’s Medium (IMDM) (Wako Pure Chemical Industries, Osaka, Japan) supplemented with 10% fetal bovine serum and 1× antibiotic-antimycotic (Sigma-Aldrich, St. Louis, MO) on culture dishes coated with rat tail type-I collagen (Corning, Corning, NY). Haploid HAP1 cells were purified by sorting based on forward scatter (FSC) intensity using a JSAN desktop cell sorter (Bay bioscience, Kobe, Japan). For each sorting, ∼1 × 10^6^ cells were collected. Sorted cells were cultured for a further 6–7 d to reach sub-confluence on 15 cm dishes (Nippon Genetics, Tokyo, Japan), and then stored in freezing medium (Bambanker, Lymphotec Inc., Tokyo, Japan) as 5–6 aliquots in vials (Corning) at -80°C or -196°C. Every cell culture lot was checked for DNA content as described below. Haploid-enriched cells were used within 7 d after recovery from frozen stocks for all experiments, to minimize the effects of spontaneous diploidization. For long-term culture experiments, the point at which haploid-enriched HAP1 cell stocks were freshly thawed was set as “day 0,” and cells were cultured for several weeks, with passaging every 1–3 d. To obtain diploid HAP1 cells, haploid-enriched cells were cultured for a few weeks, and the spontaneously diploidized cell population purified by FSC-based sorting. To obtain tetraploid HAP1 cells, diploid HAP1 cells were treated with 2.5 μM cytochalasin D for 16 h to induce cytokinesis failure, washed three times with IMDM, and then subjected to limiting dilution. After 10 d, colonies containing cells that were uniform in size and larger than diploid cells were picked and cultured, and their ploidy states were tested by DNA content analyses using flow cytometry to select tetraploid clones. The HAP1 EGFP-α-tubulin or GFP-centrin cell line was established by transfecting HAP1 cells with pEGFP-α-tubulin vector (Uehara and Goshima, 2010) or pEGFP-centrin2 vector (Kleylein-Sohn et al., 2007) (from Addgene, plasmid #41147), respectively, and selecting positive cells that grew in the presence of 500 μg/mL G418 (Wako). For DNA content analyses, HAP1 cells were cultured until they reached sub-confluence on 10 cm dishes, then trypsinized, washed once with Dulbecco’s phosphate-buffered saline (DPBS, Wako), suspended in 1 mL DPBS at a density of 2 × 10^6^ cells/mL, stained with 10 μg/ml Hoechst 33342 (Dojindo, Kumamoto, Japan) for 15 min at 37°C, washed once with DPBS, and their DNA contents analyzed using a JSAN desktop cell sorter.

### RNA interference

For siRNA transfection, Lipofectamine RNAiMAX (Thermo Fisher Scientific, Waltham, MA) was used following the manufacturer’s instructions. The siRNAs used in this study were 5 ´-CGUACGCGGAAUACUUCGATT-3 ´ (luciferase); 5 ´-CAGUUAAGCAGGUACGAAATT-3 ´ (Aug6) (Uehara et al., 2009); 5 ´-GAGUCGGGACCACAGUUUATT-3 ´ (Mad2) (Chin and Herbst, 2006); and 5 ´-UGGACGUCAUCCAAUGAGATT-3´ (PCNT) (Kim et al., 2015). Cells were subjected to live cell imaging or flow cytometry analyses 48 h after siRNA transfection.

### Immunofluorescence staining

For immunofluorescence staining (IF) of centrosomal proteins and mitotic spindles, cells or embryos were fixed with 100% methanol at -20°C for 10 min. For IF of astral MTs, cells were fixed with 3.2% paraformaldehyde (PFA) and 2% sucrose in DPBS for 10 min at 37°C, and permeabilized with 0.5% Triton-X100 in DPBS supplemented with 100 mM glycine (DPBS-G) on ice for 5 min. For IF of incorporated BrdU, cells were pre-fixed with 100% methanol at -20°C for 10 min, post-fixed with 3.7% PFA in DPBS for 15 min at 25°C, and treated with 1% Triton-X100 in 4 N HCl for 5 min at 25°C. Fixed samples were treated with BSA blocking buffer (150 mM NaCl; 10 mM Tris-HCl, pH 7.5; 5% BSA; and 0.1% Tween 20) for 30 min at 25°C, incubated with primary antibodies overnight at 4°C, and incubated with secondary antibodies for 1 h at 37°C or overnight at 4°C. Following each treatment, cells were washed 2–3 times with DPBS or DPBS-G. Stained cells were mounted with mounting medium (90% (v/v) Glycerol; 100 mM Tris-HCl, pH 8.0; and 0.5% (w/v) n-propyl gallate). Stained mouse embryos were embedded in 0.5% PrimeGel Agarose LMT (Takara Bio, Kusatsu, Japan) dissolved in DPBS.

### Immunoblotting

For immunoblotting (IB), proteins separated by SDS-PAGE were transferred to Immun-Blot PVDF membrane (Bio-Rad, Hercules, CA). Membranes were then blocked with 0.3% skim milk in TTBS (50 mM Tris, 138 mM NaCl, 2.7 mM KCl, and 0.1% Tween 20), incubated with primary antibodies overnight at 4°C or for 1 h at 37°C, and with secondary antibodies for 30 min at 37°C. Each step was followed by three washes with TTBS. Signal detection employed the ezWestLumi plus ECL Substrate (ATTO, Tokyo, Japan) and a LuminoGraph II chemiluminescent imaging system (ATTO).

### Antibodies

Antibodies were purchased from suppliers, and used at dilutions, as follows: rat monoclonal anti-α-tubulin (YOL1/34, EMD Millipore, Temecula, CA; 1:1000 for IF and 1:500 for IB); mouse monoclonal anti-β-tubulin (10G10, Wako; 1:1000 for IB); goat polyclonal anti-Histone H2B (sc-8650, Santa Cruz Biotechnology, Dallas, TX; 1:50 for IF); mouse monoclonal anti-centrin (20H5, EMD Millipore; 1:1000 for IF); rabbit polyclonal anti-centrin 2 (sc-27793, Santa Cruz Biotechnology; 1:50 for IF); mouse monoclonal anti-γ-tubulin (GTU88, Sigma-Aldrich; 1:200 for IF in HAP1 cells, 1:100 for IF in mouse embryos); rabbit polyclonal anti-Cep152 (ab183911, Abcam, Cambridge, United Kingdom; 1:1000 for IF); mouse monoclonal anti-Plk4 (6H5, EMD Millipore; 1:500 for IF); rabbit polyclonal anti-CP110 (A301-343A, Bethyl Laboratories, Montgomery, TX; 1:500 for IF); mouse monoclonal anti-SAS-6 (sc-81431, Santa Cruz Biotechnology; 1:50 for IF); rabbit polyclonal anti-C-Nap1 (14498-1-AP, Proteintech, Rosemont, IL; 1:200 for IF); rabbit polyclonal anti-PCNT (ab4448, Abcam; 1:200 for IB); rabbit polyclonal anti-Aug6 (1:500 for IB) (Uehara et al., 2009); mouse monoclonal anti-mad2 (2G9, Medical & Biological Laboratories, Nagoya, Japan; 1:000 for IB); rat monoclonal anti-BrdU (sc-56258, Santa Cruz Biotechnology; 1:50 for IF); and fluorescence- or horseradish peroxidase-conjugated secondaries (Jackson ImmunoResearch Laboratories, West Grove, PA; 1:1000 for IF and IB).

### Cell imaging

Fixed and living cells were observed under a TE2000 microscope (Nikon, Tokyo, Japan) equipped with a ×100 1.4 numerical aperture (NA) Plan-Apochromatic, a ×60 1.4 NA Plan-Apochromatic, or a ×40 1.3 NA Plan Fluor oil immersion objective lens (Nikon), a CSU-X1 confocal unit (Yokogawa, Tokyo, Japan), and an iXon3 electron multiplier-charge coupled device (EMCCD) camera (Andor, Belfast, United Kingdom) or an ORCA-ER CCD camera (Hamamatsu Photonics, Hamamatsu, Japan). Image acquisition was controlled by μManager software (Open Imaging). Long-term live imaging for cell cycle analyses was performed using a LCV110 microscope (Olympus, Tokyo, Japan) equipped with a ×40 0.95 NA UPLSAPO dry lens (Olympus). Since we found that 488-nm light irradiation severely interfered the progression of cell cycle and mitosis in HAP1 cells, we used bright-field microscope for long-term live imaging for cell cycle analyses (Fig. 1 and Fig. 7).

### Cell cycle synchronization

Asynchronous HAP1 cell cultures were treated with 20 ng/mL nocodazole (Wako) for 4 h, and then washed with IMDM supplemented with 10% fetal bovine serum and 1× antibiotic-antimycotic three times. Mitotic cells were dislodged from the culture dishes by tapping and then transferred to collagen-coated 8-well coverglass chambers (ZelKontakt, Noerten-Hardenberg, Germany) or 6-well dishes (Nippon Genetics), and synchronized cells sampled at each time point indicated. To arrest cells in early S phase, the nocodazole-released cells were treated with 4 mM thymidine (Wako).

### Theoretical modeling of the progression of diploidization

To theoretically assess the impact of haploid-specific mitotic defects and cell cycle delay on the stability of the haploid state in HAP1 cells, we constructed a mathematical model based on the following simple assumptions: 1) Haploid and diploid populations proliferate exponentially with characteristic doubling times corresponding to their cell cycle lengths (Fig. 1D); 2) haploid cells die or convert into diploids through mitotic death or mitotic slippage, respectively, with the observed frequencies (Fig. 1F); and 3) the proportion of cells in G1 phase remains unchanged in haploid populations throughout long-term culture. The basic time-dependent growth of a cell population can be modeled as:

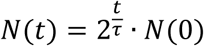

where *N* is the number of cells at time point, *t*, and τ is the average cell cycle length in the cell population.

Assuming ergodicity in the processes of the cell cycle, mitosis, and cell proliferation, the probability of the occurrence of mitotic death in a haploid cell population during unit time length, *p*_*death*_, is derived as:

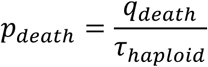

and mitotic slippage in a haploid cell population during unit time length, *p*_*slippage*_, as:

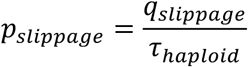

where *q*_*slippage*_ and *q*_*death*_ are the rate of incidence of mitotic slippage and mitotic death per mitotic event, respectively, and τ_haploid_ is the average cell cycle length of a haploid cell population. The time-dependent growth of a haploid cell population is then modeled as:

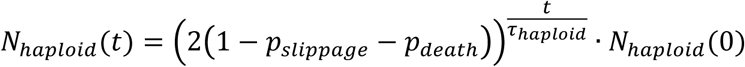

where *N*_*haploid*_ is the number of haploid cells at the time point, *t*. A haploid cell that converts to diploid through mitotic slippage at any point during the simulation will thereafter proliferate with the characteristic doubling time of diploid cells. The time-dependent growth of a diploid cell population is, therefore, modeled as:

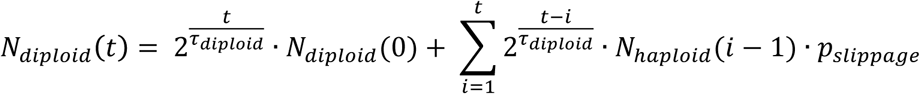

where *N*_*diploid*_ is the number of diploid cells at time point, *t*, and τ_diploid_ is the average cell cycle length of a diploid cell population. Finally, the time-dependent change in the percentage of haploid cells in G1, *P*_*haploid, G1*_, in a cell culture is modeled as:

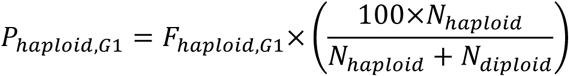

where *F*_*haploid, G1*_ is the fraction of cells in G1 phase within a haploid cell population.

Computer programs were written using MATLAB (Mathworks, Natick, MA). Parameters used in simulations are listed in Table S1.

### Image analyses

To estimate the sizes of HAP1 cells with different ploidy levels, the areas of trypsinized round-shaped cells were measured using the ROI tool in ImageJ software (National Institutes of Health, Bethesda, MD), and the cell radius, *r*, was calculated using the equation:

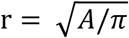

where *A* is the measured cell area.

The fluorescence intensities of centrosome-associated immunofluorescence signals were measured using plot profile or round-shaped ROIs with diameters of 0.53 μm (ImageJ), with subtraction of back-ground signals in the areas outside the cells or the cytoplasmic areas, respectively. Measurement of inter-centriolar distance in GFP-centrin cells or counting of astral fibers in fixed anaphase cells was performed using the Line tool or the Manual tracking tool in ImageJ, respectively.

### Parthenogenesis, and embryo culture

C57BL/6 and DBA/2 mice were purchased from Japan SLC, Inc. (Hamamatsu, Japan) and CBA mice were from Charles River Laboratories Japan, Inc. (Yokohama, Japan). For mouse embryo experiments, 8–12-week-old female B6D2F1 (C57BL/6 × DBA/2) or BCF1 mice (C57BL/6 × CBA) were injected with 5 IU pregnant mare serum gonadotropin (PMSG, ASKA Animal Health, Tokyo, Japan) followed by injection with 5 IU human chorionic gonadotropin (hCG, ASKA Pharmaceutical, Tokyo, Japan) 46–48 h later, and matured oocytes were obtained from oviducts 16 h later. Haploid and diploid parthenogenetic embryos were produced as described previously with slight modifications (Kishigami and Wakayama, 2007; Latham et al., 2002). Oocytes were treated with 0.1% hyaluronidase (Sigma-Aldrich) in M2 medium (Sigma-Aldrich) for 30 s to remove cumulus cells, washed with M2 medium and M16 medium (Sigma-Aldrich) three times each, and incubated in M16 medium supplemented with 2 mM EGTA (EGTA-M16) for 20 min. To produce haploid parthenogenetic embryos, these oocytes were treated with 5 mM SrCl_2_ in EGTA-M16 for 2.5 h. Diploid parthenogenetic embryos were produced by treating the oocytes with 5 mM SrCl_2_ in EGTA-M16 in the presence of 5 μg/mL cytochalasin B (Wako) for 2.5 h, and then incubating them in KSOM medium (MTI-GlobalStem, Gaithersburg, MD) in the presence of the same concentration of cytochalasin B for 3.5 h. Activated haploid and diploid parthenogenetic embryos were then washed with KSOM three times and cultured in the same medium at 37°C with 5% CO_2_.

### Ethics statement

The maintenance and handling of mice for all embryo experiments were performed in the animal facility of the Platform for Research on Biofunctional Molecules of Hokkaido University under the guidelines and with the permission of the committee on animal experiments of Hokkaido University (permission number 16-0038).

## Acknowledgments

We are grateful to Mithilesh Mishra, Sarada Bulchand, Tomoko Kamasaki, and Gohta Goshima for valuable comments on the draft of this paper, Meng-Fu Bryan Tsou for GFP-centrin live imaging advice, Kuniharu Ijiro, Shin-Ichiro Nishimura, Min Yao, Jian Ping Gong, and other members of the Faculty of Advanced Life Science and the Open Facility (Hokkaido Univ.) for the use of their equipment. This work was supported by the Akiyama Life Science Foundation, the Inamori Foundation, the Mochida Memorial Foundation, the Naito Foundation, the Nakajima Foundation, the Noastec Foundation, the SGH Foundation, the Suhara Memorial Foundation, the Sumitomo Foundation, the Takeda Science Foundation, the Uehara Memorial Foundation, and MEXT/JSPS KAKENHI (Grants Numbers 15K14501, and 17K15111) to R. U. The authors declare no competing financial interests.

## Author Contributions

Conceptualization, K.Y., and R.U.; Methodology, K.Y., R.M., A.S., K.K., T.K., and R.U.; Software, R.M., Y.T., and R.U.; Investigation, K.Y., R.M., T.Y., and R.U.; Formal Analysis, K.Y., R.M., T.Y., and R.U.; Resources, T.K., and R.U.; Writing – Original Draft, R.U.; Writing – Review & Editing, Y.T., T.K., and R.U.; Funding Acquisition, T.K., and R.U.

---

**Fig. S1:**
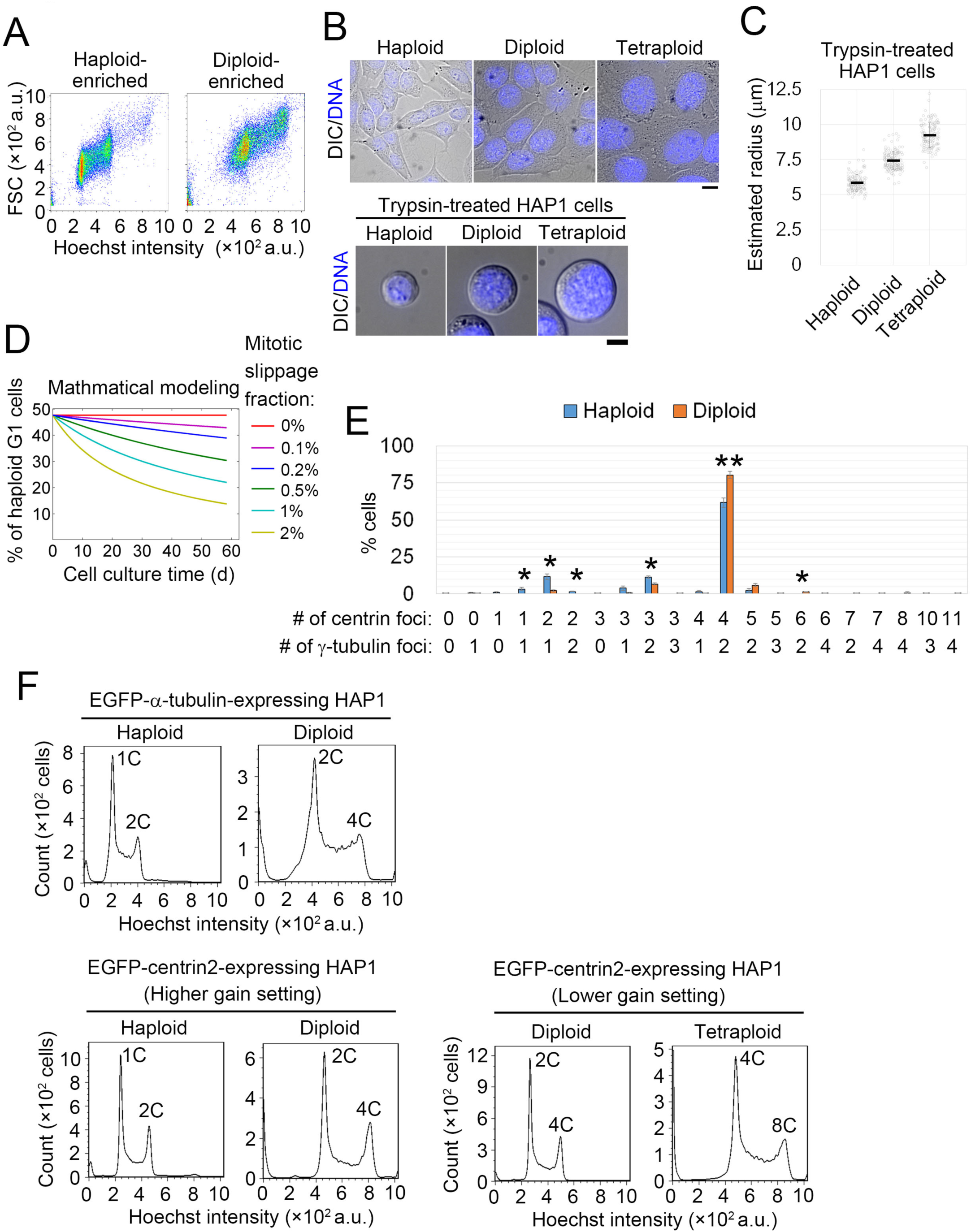
Analysis of isogenic HAP1 cell lines with different ploidies. (**A**) Flow cytometric analysis of FSC and DNA content in Hoechst-stained haploid or diploid HAP1 cells. (**B**) Microscopy of adhesive or trypsin-treated haploid, diploid, or tetraploid HAP1 cells. DNA was stained using Hoechst. Scale bars, 10 μm. (**C**) Radii of trypsin-treated haploid, diploid, or tetraploid HAP1 cells, estimated using measured values of cell area. Means ± SE of at least 125 cells from 2 independent experiments per condition. There were significant differences between all pairs of samples (p < 10^-50^, *t*-test). (**D**) Time plots of simulated haploid G1 fractions during long term culture of haploid enriched cell populations with different frequencies of mitotic slippage. Note that neither haploid-specific cell cycle delay nor mitotic death was taken into account in these simulations. (**E**) Distribution of centriole and centrosome number in mitotic haploid and diploid HAP1 cells shown in Fig. 2D. Means ± SE of 3 independent experiments. Statistically significant differences between haploid and diploid cells are indicated by asterisks (* p < 0.05, ** p < 0.01, *t*-test). At least 302 cells were analyzed per condition. (**F**) Flow cytometric analysis of DNA content in Hoechst-stained haploid or diploid HAP1 cells stably expressing EGFP-α-tubulin or GFP-centrin. GFP-centrin lines were analyzed using two different gain settings.

**Fig. S2:**
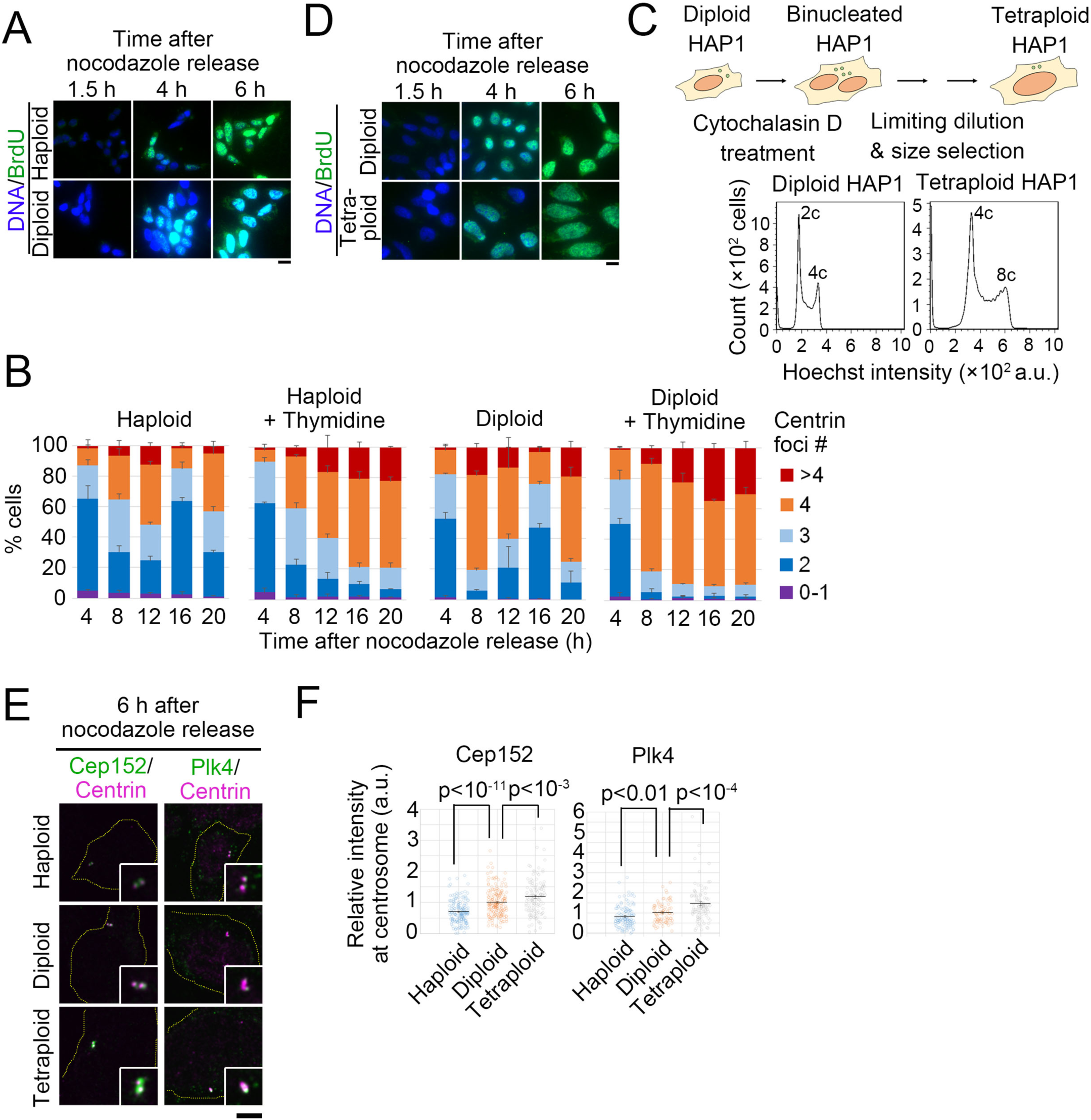
Ploidy-dependent changes in centrosome duplication efficiency. (**A, D**) Immunostaining of incorporated BrdU in synchronized HAP1 cells with different ploidies. DNA was stained with DAPI. Scale bars, 10 μm. (**B**) Percentages of haploid or diploid cells with indicated numbers of centrin foci at each time point after nocodazole release. Cells were incubated with or without 4 mM thymidine. Mean ± SE of 3 independent experiments. At least 161 cells were analyzed for each data point. (**C**) A schematic of the procedure for the establishment of tetraploid HAP1 cells (top), and flow cytometric analysis of DNA content in Hoechst-stained diploid or tetraploid HAP1 cells (bottom). (**E**) Immunostaining of centrosome proteins in synchronized cells with different ploidies. Broken lines indicate boundaries of cells or nuclei. Insets: ×2.5 enlarged images of the centrioles. Scale bar, 5 μm. (**F**) Normalized intensities of centrosome proteins at the centrosomes in E. Mean ± SE of at least 65 centrioles from 2 independent experiments for each condition (*p* values calculated using *t*-tests are shown).

**Fig. S3:**
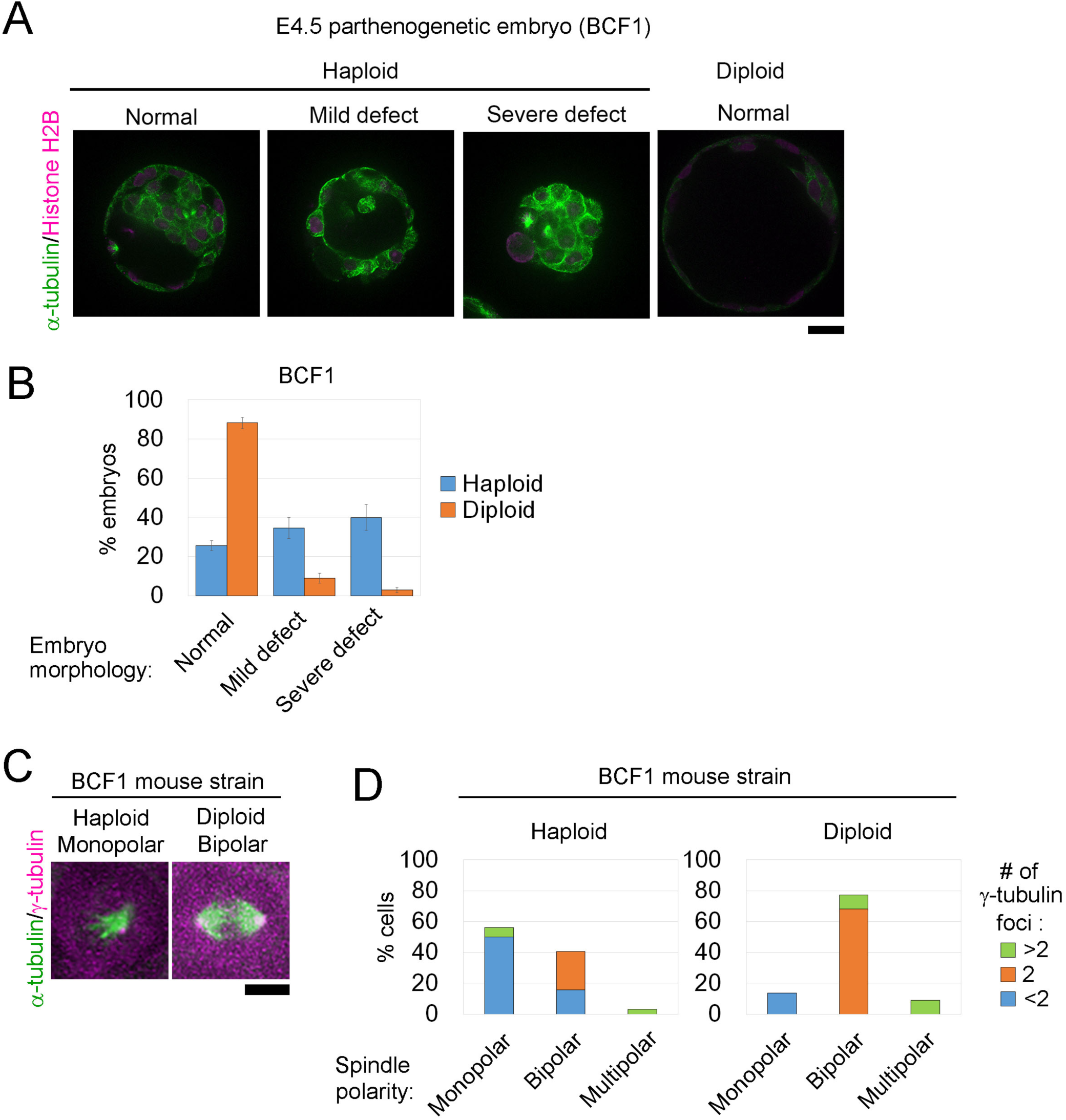
Centrosome reduction and monopolar spindle formation in mouse haploid embryos. (**A, C**) Immunostaining of α-tubulin and histone H2B (A), or γ-tubulin (C) in haploid and diploid mouse parthenogenetic embryos (stage E4.5) from BCF1 strain. (C) Enlarged mitotic cells. Scale bars: 20 μm in A, and 5 μm in C. (**B**) Frequencies of morphological abnormalities in A. Means ± SE of 3 independent experiments. At least 71 embryos were analyzed per condition. (**D**) Frequency of spindle polarities and centrosome numbers in C. At least 22 cells from 18 embryos from 3 independent experiments were analyzed per condition.

## Video 1

Live imaging of haploid HAP1 cells that completed cell division (blue arrow). Pink arrows indicate daughter cells. Scale bar, 5 μm.

## Video 2

Live imaging of haploid HAP1 cells that underwent mitotic slippage (blue arrow). Scale bar, 5 μm.

## Video 3

Live imaging of diploid HAP1 cells that completed cell division. Scale bar, 5 μm.

## Video 4

Live imaging of haploid HAP1 EGFP-α-tubulin cells with bipolar spindles (blue arrow). Pink arrows indicate daughter cells. Scale bar, 5 μm.

## Video 5

Live imaging of haploid HAP1 EGFP-α-tubulin cells, in which monopolar spindles later converted to bipolar (blue arrow). Pink arrows indicate daughter cells. Scale bar, 5 μm.

## Video 6

Live imaging of haploid HAP1 EGFP-α-tubulin cells that underwent mitotic slippage (blue arrow). Scale bar, 5 μm.

## Video 7

Examples of astral MT tracing for the quantification in Fig. 9B. Scale bar, 5 μm.

## Video 8

Examples of astral MT tracing for the quantification in Fig. 9F. Scale bar, 5 μm.

---

**Table S1:** Parameters and constants used in the simulations

| Constants<br>(Unit) | Meaning | Fig. 1G<br>w/ cell cycle<br>delay,<br>mitotic<br>slippage,<br>and mitotic<br>death | Fig. 1G<br>w/ cell cycle<br>delay, and<br>mitotic<br>death | Fig. 1G<br>w/ mitotic<br>slippage | Fig. S1D |
| --- | --- | --- | --- | --- | --- |
| $N_{haploid}(0)$<br>($\times 10^3$ cells) | Initial<br>number of<br>haploid cells | 5 | 5 | 5 | 5 |
| $N_{diploid}(0)$<br>($\times 10^3$ cells) | Initial<br>number of<br>diploid cells | 0 | 0 | 0 | 0 |
| $q_{slippage}$<br>(h) | Incidence<br>rate of<br>mitotic<br>slippage per<br>mitotic<br>event | 0.59 | 0 | 0.59 | 0-2 |
| $q_{death}$<br>(h) | Incidence<br>rate of<br>mitotic<br>death per<br>mitotic<br>event | 1.6 | 1.6 | 0 | 0 |
| $\tau_{haploid}$<br>(h) | Cell cycle<br>length of<br>haploid | 13.4 | 13.4 | 11.9 | 11.9 |
| $\tau_{diploid}$<br>(h) | Cell cycle<br>length of<br>diploid | 11.9 | 11.9 | 11.9 | 11.9 |
| $F_{haploid, G1}$<br>(%) | G1 cell<br>fraction<br>within<br>haploid<br>population | 47.7 | 47.7 | 47.7 | 47.7 |

